# A disease-linked lncRNA mutation in RNase MRP inhibits ribosome synthesis

**DOI:** 10.1101/2021.03.29.437572

**Authors:** Nic Robertson, Vadim Shchepachev, David Wright, Tomasz W. Turowski, Christos Spanos, Aleksandra Helwak, Rose Zamoyska, David Tollervey

**Affiliations:** Wellcome Centre for Cell Biology, University of Edinburgh, Edinburgh, UK; Ashworth Laboratories, Institute of Immunology and Infection Research, University of Edinburgh, Edinburgh, UK

**Keywords:** protein-RNA interaction, RNA-binding sites, UV crosslinking, mass spectrometry, genetic disease, Cartilage Hair Hypoplasia, ribosome synthesis, T cell activation

## Abstract

*RMRP* encodes a non-coding RNA forming the core of the RNase MRP ribonucleoprotein complex. Mutations cause Cartilage Hair Hypoplasia (CHH), characterized by skeletal abnormalities and impaired T cell activation. Yeast RNase MRP cleaves a specific site in the pre-ribosomal RNA (pre-rRNA) during ribosome synthesis. CRISPR-mediated disruption of *RMRP* in human cells lines caused growth arrest, with pre-rRNA accumulation. Here, we analyzed disease-relevant primary cells, showing that mutations in *RMRP* impair mouse T cell activation and delay pre-rRNA processing. Patient-derived human fibroblasts with CHH-linked mutations showed similar pre-rRNA processing delay. Human cells engineered with the most common CHH mutation (70^AG^ in *RMRP*) show specifically impaired pre-rRNA processing, resulting in reduced mature rRNA and a reduced ratio of cytosolic to mitochondrial ribosomes. Moreover, the 70^AG^ mutation caused a reduction in intact RNase MRP complexes. Together, these results indicate that CHH is a ribosomopathy, and the first processing-specific human disorder to be described.

**Highlights:**

- Mutations in *RMRP* lncRNA impair pre-rRNA processing and T cell activation
- Patient derived fibroblasts show impaired pre-rRNA processing
- Cells with the most common disease-linked mutation have specific processing defects
- Cytoplasmic ribosomes and intact RNase MRP complexes are also reduced in these cells

## Introduction

*RMRP* mutations are associated with a spectrum of disorders presenting with skeletal dysplasia, abnormal hair and immune deficiency with impaired T-cell activation^1^. The most common of these syndromes is called Cartilage Hair Hypoplasia (CHH). Patients with CHH have reduced life expectancy due to immune deficiency, but the mechanism underlying this problem is not understood^1^. Clinically, patients often experience recurrent infections, and also have a higher incidence of auto-immunity and cancer. At a cellular level, lymphocytes from CHH patients show reduced proliferation in response to stimulation and increased activation-induced cell death^2,3^. Naïve T cells are small and quiescent, but on activation increase in size over 24h before beginning to rapidly divide^4,5^. This activation program is accompanied by a 10-fold increase in per-cell ribosome abundance, achieved through upregulation of both ribosomal protein (RP) and ribosomal RNA (rRNA) production^6,7^.

The human *RMRP* gene encodes a non-coding RNA, which associates with around 10 proteins in the RNase MRP complex^8,9^. RNase MRP was initially proposed to function in cleavage of an RNA primer during mouse mitochondrial DNA replication, giving its name: <u>m</u>itochondrial <u>R</u>NA <u>p</u>rocessing^9,10^. However, in budding yeast, RNase MRP is required for cleavage at a specific site in the pre-ribosomal RNA^8,11,12^. *RMRP* is an essential gene in all studied eukaryotes, but a role, if any, in mammalian ribosome synthesis was unclear^13^. However, CRISPR-mediated disruption of *RMRP* in human immortalized cell lines was recently reported to cause growth arrest, with an accumulation of ribosomal precursor RNA (pre-rRNA)^14^. The evolutionarily-related ncRNA *RPPH1* forms the core of the RNase P complex, which processes pre-tRNAs and shares multiple protein components with RNase MRP.^9^

Human rRNAs are transcribed as a long precursor, 47S pre-rRNA. This is processed in a complex series of reactions (Fig. S1), to the 18S rRNA, destined for the small ribosomal subunit (SSU), and 28S and 5.8S, present in the large subunit (LSU)^15^. A third LSU rRNA, 5S, is transcribed separately. By analogy with yeast, RNase MRP is thought to mediate cleavage at site 2 within Internal Transcribed Spacer 1 (ITS1) that separates precursors to the 18S and 5.8S/28S rRNAs^14^. Whether disease-causing mutations act through disrupted ribosome synthesis has not been addressed.

In this study we show that mutations in *RMRP* impair mouse T-cell activation and delay pre-ribosomal RNA (rRNA) processing, phenotypes recapitulated in patient-derived human fibroblasts and in cells engineered with the most common CHH mutation.

## Methods

### CRISPR targeting of RMRP in primary mouse T cells

Rag1 KO, C57BL6/J mice were bred at the University of Edinburgh under a project license granted by the UK Home Office and following institutional ethical guidance. Peripheral lymph nodes were dissected from Rag1 knockout mice homozygous for the OT1 allele. Obtained cells were stained with a division tracker dye (CellTrace Violet; Invitrogen; cat. C34557), then activated at 250,000 cells/mL with 10 nM N4 peptide (peptide sequence SIINFEKL). After 24 hours, cells were transfected with Cas9 protein and guide RNAs targeting *Rmrp*, using a Neon Transfection System (Invitrogen). Cells were then cultured for a further 48 hours in media containing recombinant IL-2 (20 ng/mL). For ICE analysis, an amplicon spanning *Rmrp* was sequenced, and traces analysed using the Synthego Inference of CRISPR Edits (ICE) tool web interface^16^.

### Human cell culture

K562 cells were obtained from ATCC (cat. CCL-243), and grown in RPMI 1640 medium with 10% FBS. Patient and control fibroblasts were obtained from the Great North Biobank (REC number 16-NE-0002), and grown in DMEM with 10% FBS.

### Northern Blotting

RNA was extracted using Trizol (Invitrogen; cat. 15596026), and resolved on either an 8% acrylamide gel with 8.3 M urea or a 3% agarose gel. RNA was then transferred to a nylon membrane for probing using oligonucleotide probes labelled with. ^32^P-*γ*ATP.

### Generation of stable CRISPR-edited human cell lines

Parental cells were nucleofected with mix including guide RNA, Cas9 or Cpf1 protein, and synthesised, single stranded repair templates including the intended tag or mutation flanked by homology arms. Clones obtained from single cells were screened by PCR and sequencing after two weeks.

### FlowFISH

Cells were fixed, permeabilised, and stained with fluorescently-labelled oligonucleotide probes complementary to 18S or 28S rRNA before analysis by flow cytometry, as described.^17^

### Total RNA-Associated Protein Purification (TRAPP)

Cells were grown for 10 divisions in SILAC RPMI (Thermo Fischer; cat. 88365) supplemented with 50 µg/L each of lysine and arginine. For “light” cultures, these amino acids were obtained from Sigma. For “heavy” cultures, ^13^C_6_-lysine and ^13^C_6_-arginine were obtained from Cambridge Isotope Laboratories (cat. CLM-226 and CLM-2247, respectively). Cells were grown to a density of 0.5 - 0.8 × 10^6^ cells / mL, then cross-linked with 400 mJ/cm^2^ of UVC using a Vari-X-Link device^18^. Heavy and light samples were then mixed 1:1 based on nucleic acid content, and RNA-associated proteins purified following a modified version of the published TRAPP protocol, using silica columns in place of silica beads^19^. Proteins were digested on the column with 0.25 µg of Trypsin/Lys-C protease mix (Promega; cat. V5071), and peptides eluted for mass spectrometry.

### Cross-linking and analysis of cDNAs (CRAC)

For yeast CRAC, standard methods were used to insert a HIS-TEV-Protein A (HTP) tag to the genomic copy of Pop1. CRAC was performed following a published protocol^20^. For human cell CRAC, samples were cross-linked as for TRAPP, except that normal media was used rather than SILAC media. Human CRAC followed the yeast CRAC protocol with minor technical modifications. Data were analysed using custom scripts calling tools from the PyCRAC collection^21^.

### Statistics

All statistics were calculated using Prism 9.0.0, with tests as indicated in figure legends. In general, plots show mean values +/- standard deviations.

## Results

### Disruption of *RMRP* impairs T cell proliferation and rRNA processing

Activating T cells require high rates of ribosome synthesis to support rapid cell division^6^. We hypothesized that this might confer particular vulnerability to partial disruption of *RMRP* function if its primary biological role is in pre-rRNA processing. To test this, we used T cells from OT-1 Rag^-/-^ mice, all of which recognize the same ovalbumin-derived antigen. Naïve primary transgenic T cells were isolated, stained with a division tracker dye (CellTrace Violet), and activated by culturing with ovalbumin peptide (Fig. 1A). After 24h, we disrupted *RMRP* by electroporating these cells with CRISPR components, including one of four guides targeting *RMRP*, or a no-guide control (mock). Cells were then left to proliferate for a further 48h. At this time point, guide efficiency was estimated by calculating the proportion of alleles in the population containing an indel, using the Synthego ICE tool to deconvolute Sanger sequencing traces obtained from bulk populations^16^. This showed that guides 1 and 2 disrupted about 60% of alleles, whereas guides 3 and 4 were less efficient, mutating approximately 20% and 10%, respectively (Fig. 1B). No mutations were detected in the control. Proliferation after 48h was assessed by flow cytometry using a division tracker dye, CellTrace Violet, which halves in fluorescent intensity each division (Figs. 1C and 1D). This showed that disruption of *RMRP* markedly impaired T cell proliferation. Guide 1 again had the most profound effect, reducing expansion of the culture to about 60% of that seen in mock-transfected cells. Guide 3 reduced expansion to around 70% of mock-treated cultures, while guides 2 and 4 showed a modest effect (about 80%). These analyses were restricted to viable cells, as assessed by a dye for live/dead cells (Zombie Red), implying that non-lethal mutations in *RMRP* slow T cell division.

**Figure 1:**
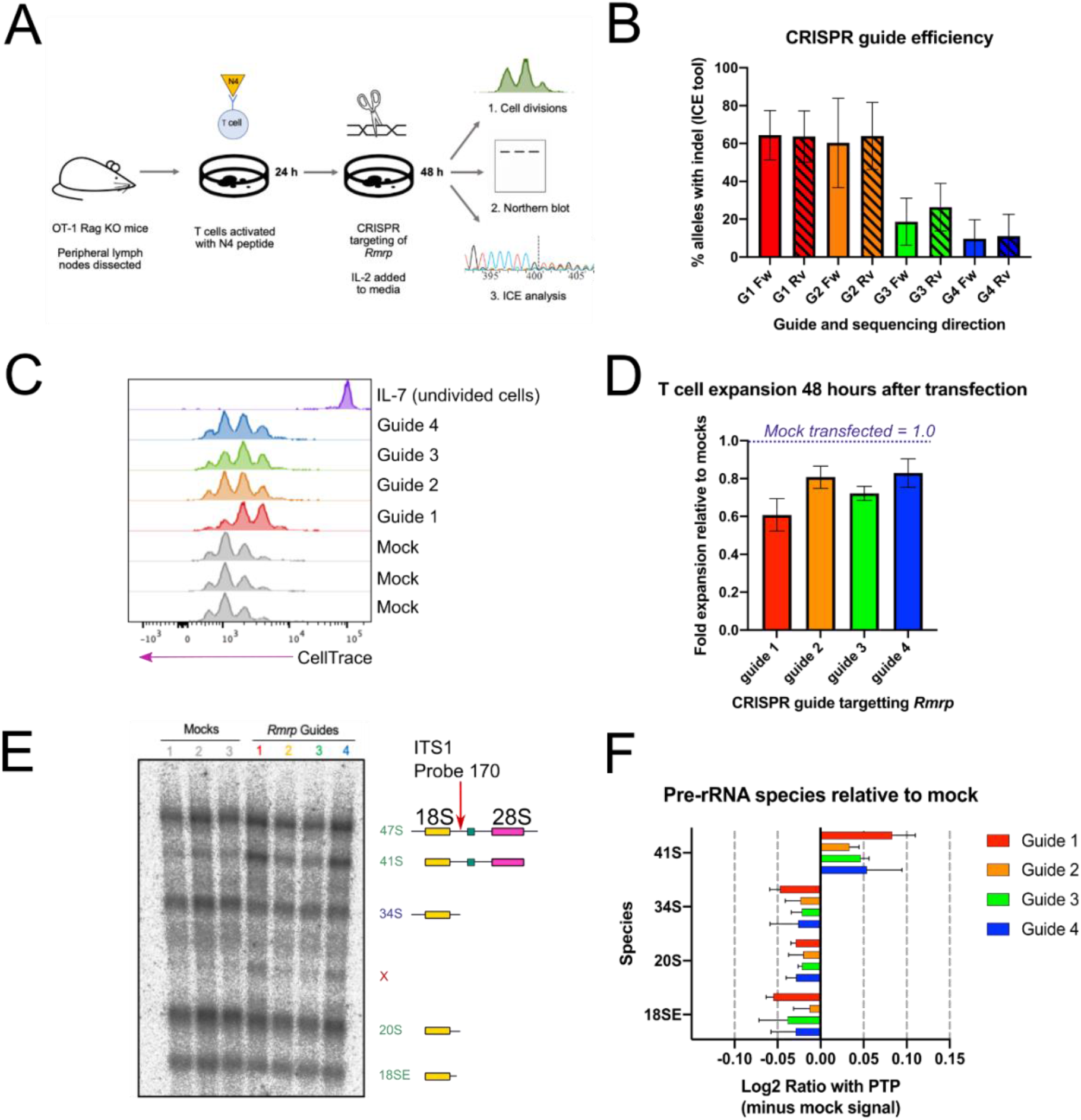
Disruption of *RMRP* impairs T cell proliferation and rRNA processing. A: Outline of experiment. Naïve TCR transgenic T cells were obtained from mouse peripheral lymph nodes and activated with N4 peptide. After 24 hours, cells were electroporated with a CRISPR mix targeting *RMRP*, or a mix without the guide (mock transfected controls). After 48 hours, cells were analyzed by flow cytometry for proliferation, Northern blotting for pre-rRNA species, and ICE analysis to determine CRISPR guide efficiency. B: CRISPR guide efficiency as determined by the Synthego Inference of CRISPR Edits (ICE) tool. A 669 bp region including *RMRP* was amplified from DNA extracted from CRISPR-targeted cultures, and sequenced with forward (Fw) and reverse (Rv) primers. The ICE tool deconvolutes Sanger sequencing traces to estimate the proportion of alleles in the population containing an indel. Graph summarizes data from three independent experiments. C: T cell proliferation 48 hours after CRISPR-targeting of *RMRP*, as measured with division tracker dye CellTrace Violet. Gated on live cells, as shown in Fig. S2A. D: Expansion of T cell cultures, calculated from data illustrated in panel C. Includes data from three independent experiments. In each replicate, expansion of CRISPR-targeted cultures was normalized to the average of mock-targeted controls in that experiment. E: Pre-rRNA species detected by a probe in ITS1, in RNA from T cells 48 hours after disruption of *RMRP*. Cartoon on right shows species represented by each band. Pre-rRNA species indicated in green are present in both Pathway 1 and Pathway 2 (Fig. S1). 41S in brown is a Pathway 1 intermediate, and those in blue are Pathway 2 species. F: Quantification of pre-rRNA species shown in E. X indicates a non-canonical precursor; analogous bands were seen in human cells with non-specifically disrupted *RMRP*^14^. Intensity values for each band were first normalized to the co-running 47S/45S band (primary transcript-plus, PTP) in that lane. For each rRNA species, the average intensity signal from the mock controls in that experiment was then subtracted. The resulting value was Log2 transformed. Panels B, D and F show mean values from three independent experiments, with SD.

We next assessed whether these growth defects are associated with impaired rRNA processing. RNA was extracted from cultures 48h after transfection and analyzed by Northern blotting using pre-rRNA probe 170 (Fig. 1E, left). This probe hybridizes in ITS1 5’ to the site 2, thus detecting pre-rRNAs that have not undergone cleavage at site 2, potentially by RNase MRP (Fig. 1E, right cartoon, and Fig. S1)^15,22^. For quantitation, pre-rRNAs levels were analyzed by the Ratio Analysis of Multiple Precursor (RAMP) method^23^, in which the abundance of each pre-rRNA species is compared to the abundance of the 47S primary transcript plus the 45S pre-rRNA, which are not resolved (designated <u>p</u>rimary transcript <u>p</u>lus; PTP; Fig. 1F). All cultures treated with *RMRP* guides showed elevated levels of 41S pre-rRNA, the intermediate immediately downstream of 47S/45S, whereas later intermediates were depleted. These results indicate that removal of the 5’ external transcribed spacer (5’ ETS) was unaffected, but cleavage in ITS1 to separate the LSU and SSU precursors is delayed (Fig. 1S).

The pre-rRNA processing defect was most prominent after targeting with guide 1, the guide that had shown the greatest effect on proliferation and produced the largest proportion of mutated loci. However, the effect of other guides did not correlate well with their effect on proliferation. Guides 2 and 4 had similar effects on proliferation, but guide 4 caused more 41S accumulation than guide 2.

Overall, these results support the model that disruption of *RMRP* impairs T cell proliferation by slowing pre-rRNA processing. However, disruption of different sites in the *RMRP* gene had different relative effects on cell division and pre-rRNA accumulation. We speculate that some guides preferentially cause lethal mutations in a smaller fraction of cells, while others induce milder defects in a larger population.

### Mutations associated with Cartilage Hair Hypoplasia impair pre-rRNA processing

Fibroblasts from CHH patients have reduced growth rates and cell cycle defects^24^. We tested whether these problems are associated with impaired pre-rRNA processing, by measuring pre-rRNA species in patient-derived, CHH fibroblasts (Fig. 2A). Cells from healthy volunteers were used as controls. Being slow growing, these cells contained substantially less pre-rRNA than activated T cells, so obtaining high quality RNA quantitation was challenging. Unlike *RMRP*-disrupted mouse T cells, the 41S pre-rRNA in patient fibroblasts was not clearly increased relative to the 47S/45S band (Fig. 2B). 41S pre-rRNA is generated in processing Pathway 1, by initial removal of both the 5’ ETS and 3’ ETS from 47S, leaving the SSU and LSU rRNA precursors joined (Fig. S1)^22^. In Pathway 2, initial cleavage at site 2 in ITS1 separates the 30S (precursor for the SSU) and 32S pre-rRNA (precursor for the LSU). Relative to 41S, the 30S pre-rRNA was significantly less abundant in CHH fibroblasts than controls. We propose that slowed site 2 cleavage, caused by reduced RNase MRP activity, leads to preferential utilization of Pathway 1 over Pathway 2 in CHH patient cells (Fig. 2B).

**Figure 2:**
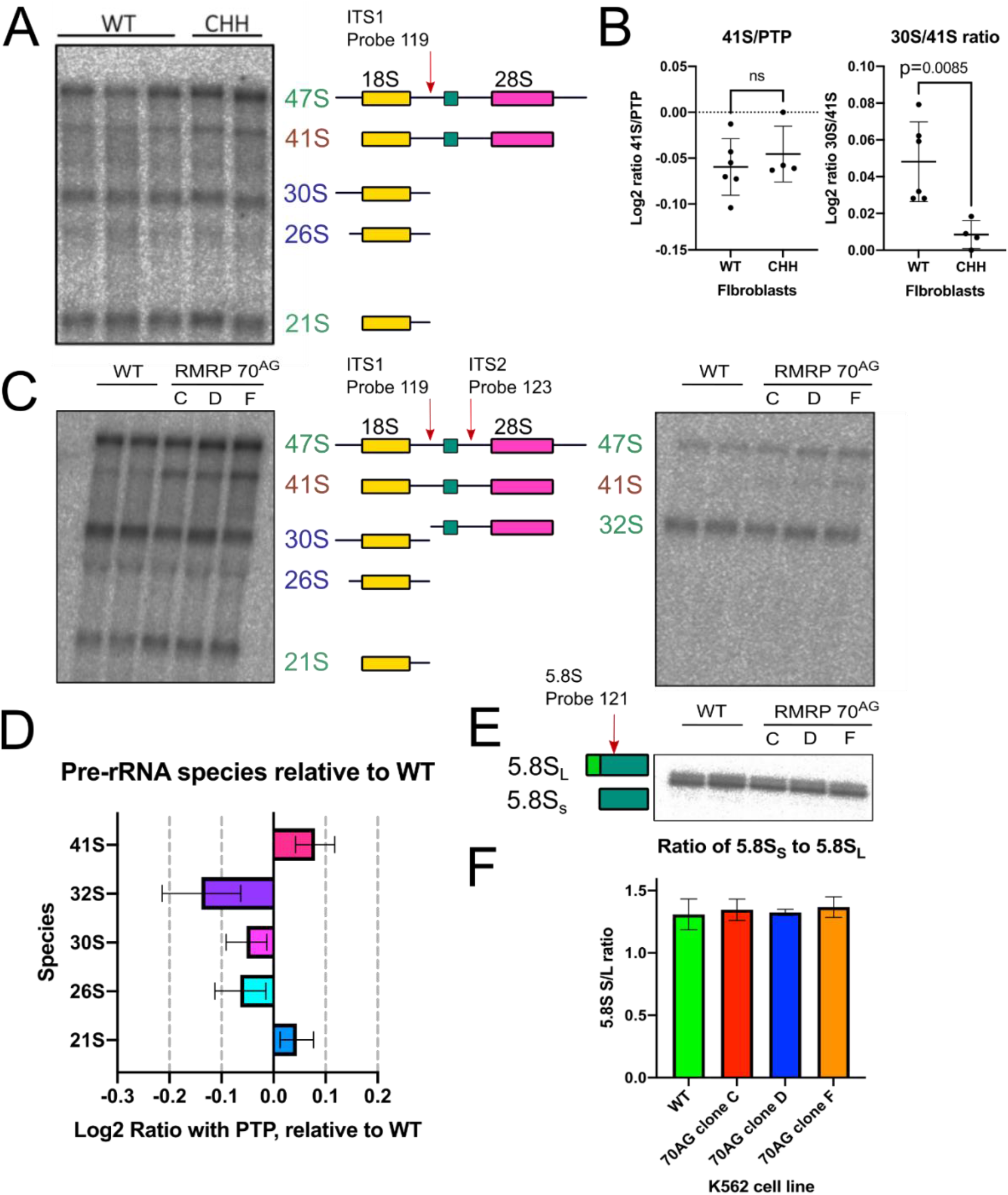
Mutations associated with Cartilage Hair Hypoplasia impair rRNA processing. A: Northern blot using ITS1 probe (probe 119) to detect pre-rRNA species in RNA extracted from CHH or healthy control (WT) fibroblasts. Pre-rRNA species indicated in green are present in both Pathway 1 and Pathway 2. 41S in brown is a Pathway 1 intermediate, and those in blue are Pathway 2 species. B: Quantification of relative Each dot represents an independent sample, and lines show mean +/- SD. Includes samples processed in two independent Northern blotting experiments. Indicated p value for 30/41S derived from two-tailed t test (t=3.463, df=8). C. Northern blot using probes against ITS1 (left; probe 119) and ITS2 (right; probe 123), to detect pre-rRNA in parental K562 cells, or CRISPR-generated clones of K562 with a 70^AG^ mutation in *RMRP*. C, D and F represent cell lines derived from independent CRISPR clones. D: Quantification of pre-rRNA species from WT and 70^AG^ K562 cells. Intensity of bands for each indicated species were first normalized to the combined 47S/45S band (PTP) in that lane. Values from WT cells were then subtracted from mutant values. Includes data from three independent experiments. E: Northern blot with probe against 5.8S rRNA, showing the short (5.8S_S_) and long (5.8S_L_) form of these species in indicated cell lines. F: Quantification of ratio between 5.8SS and 5.8SL, determined from Northern blots illustrated in E. Includes data from three independent rounds of RNA extraction, run on two separate gels. Panels D and F show mean values from three experiments, with SD.

To overcome the challenges of working with pre-RNA from fibroblasts, we generated a cell line homozygous for the most common CHH-associated mutation, an A to G transition at position 71 on the current reference sequence (NBCI sequence NR_003051.3; Fig. S3A)^1^. Prior literature refers to this mutation as 70^AG^ based on previous reference sequences, and we use 70^AG^ for consistency. K562 cells, a suspension line derived from chronic myelogenous leukemia, were selected as the parental line^25^. The 70^A^ site overlaps a PAM site compatible with CRISPR-Cas12a (previously Cpf1), allowing the 70^AG^ mutation to be introduced without creating extraneous mutations in the ncRNA^26^. Four independent, homozygous CRISPR clones were obtained (Fig. S3B). Levels of the *RMRP* ncRNA were not significantly reduced in these cells compared to wildtype levels, when assessed by qPCR (Fig. S3C). For unknown reasons, one clone (clone C) showed slightly reduced levels of the ncRNA *RPPH1*, which forms the related RNase P complex (Fig. S3C)^9^. All tested 70^AG^ clones grew on average more slowly than parental cells: over 48h, mutant cell cultures expanded 25% less than wildtype controls (Fig. S3D).

The cell lines gave good yields of high-quality RNA for pre-rRNA Northern blots (Fig. 2C). RAMP analyses with an ITS1 probe (Probe 119; Fig. 2C left) showed that the 41S:PTP ratio was consistently increased in 70^AG^ mutants, with an average band intensity 0.08 (Log2) above wildtype. In contrast, the ratios of 30S:PTP and 26S:PTP were reduced, with an average change of -0.05 and -0.06, respectively (Fig. 2D). An ITS2 probe (Probe 123; Fig. 2C right), showed a visible reduction in 32S relative to the PTP band. Quantifying this showed that 32S:PTP was decreased by an average of -0.15 fold. We conclude that more 47S is processed in Pathway 1 (via 41S) in 70^AG^ mutants, with a decrease in Pathway 2 species (notably 30S and 26S), as found in CHH patient cells.

Mature 5.8S rRNA is present in two different forms in all studied animals, fungi and plants^27-29^. In *S*.*cerevisiae* about 80% of 5.8S is the short form (5.8S_S_), which is generated by 5’ exonuclease activity following MRP cleavage^11,12,27^. An alternative, MRP-independent pathway generates 5.8S_L_, which is 5’ extended into ITS1 by 7 or 8 nucleotides. In consequence, loss of MRP activity in yeast results in a reduction in 5.8S_S_ relative to 5.8S_L_.

It was previously reported that fibroblasts from CHH patients have perturbations in the 5.8S rRNA population, with an increased abundance of species with a 5’ extension into ITS1, as measured by qPCR^30^. We assessed the ratio of 5.8S_S_ to 5.8S_L_ in the 70^AG^ K562 cells, using acrylamide-gel Northern blotting to give nucleotide-level resolution (Fig. 2E). 5.8S_S_ and 5.8S_L_ were clearly resolved, with 5.8S_S_ being more abundant. However, the 5.8S_S_ to 5.8S_L_ ratio was the same in parental and 70^AG^ cells, at approximately 1.25 (Fig. 2F). It may be that the reported, extended 5.8S rRNA qPCR products were generated by accumulated pre-rRNAs, particularly 41S, which includes the ITS1-5.8S region (Fig. S1)^30^.

### The 70^AG^ mutation in *RMRP* reduces cytosolic ribosome abundance

We next assessed if the processing delay in 70^AG^ cells resulted in a lower abundance of mature rRNA species. This is difficult to quantify using Northern blotting because rRNA comprises the majority of cellular RNA^31^. Changes in rRNA abundance are masked when a set quantity of total RNA is loaded on the gel. Instead, we used flow cytometry to measure per-cell 18S and 28S RNA signals, using fluorescently-labelled oligonucleotide probes, a technique called FlowFISH (Flow-cytometry based fluorescently-labelled in-situ hybridization)^6,17^. Probes with scrambled nucleotide sequences were used as negative controls, and gave low background (Fig 3A). All 70^AG^ clones tested had significantly reduced rRNA signals compared to the parental lines (Fig. 3B), but there was some variability. Clone C showed the greatest reduction, with an rRNA signal at about 60% of that from parental cells. Clone D showed approximately 80% of wildtype signal, with clone F at 70%.

**Figure 3:**
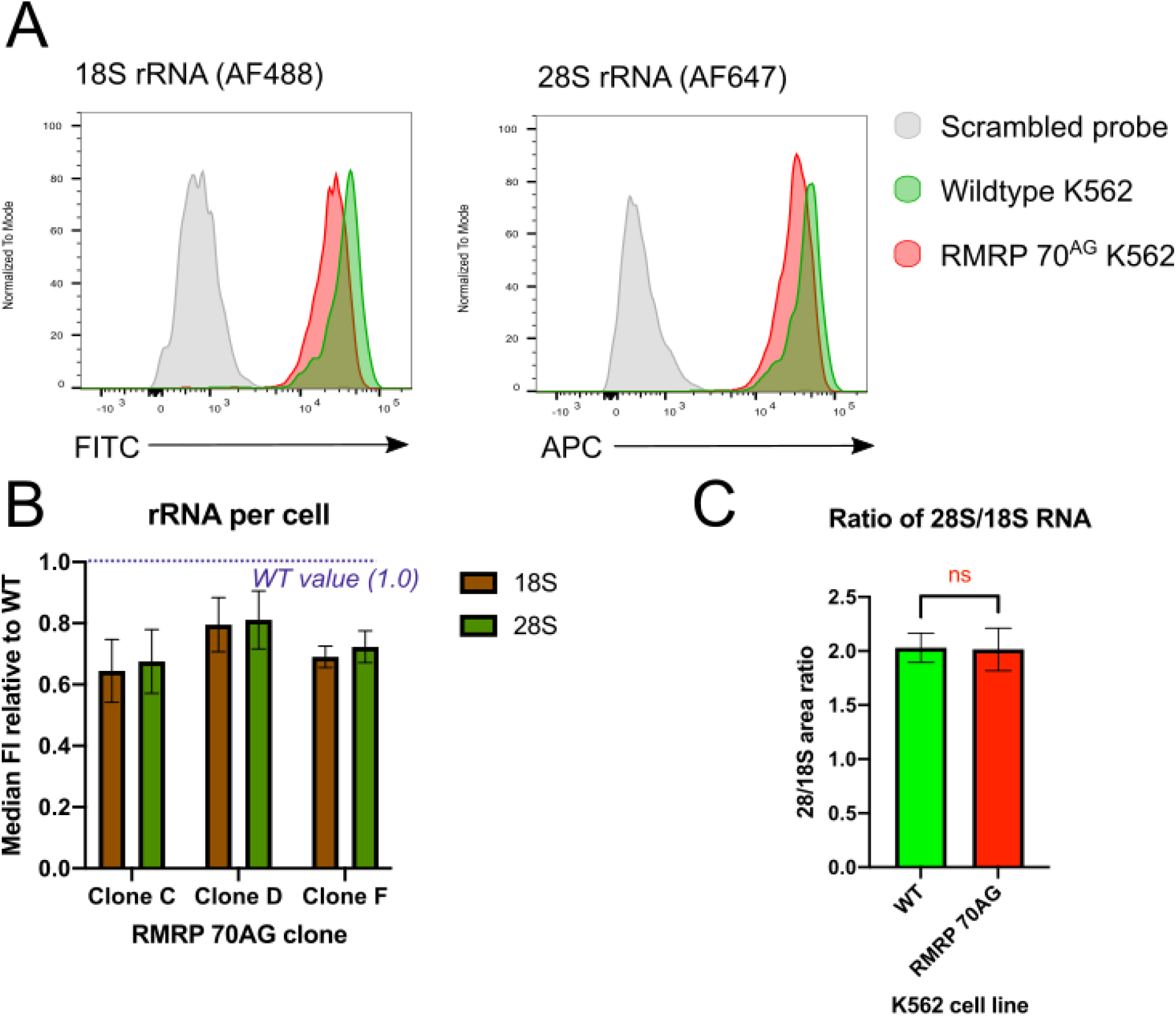
70^AG^ mutation in *RMRP* causes a reduction in rRNA per cell. A: Example data from FlowFISH experiments, showing 18S and 28S rRNA detected by fluorescently-labelled antisense oligonucleotide probes, or signal from control probes with scrambled sequences. Gated on morphologically live cells. B: Quantification of rRNA signals obtained in FlowFISH experiments. In each experiment, intensity values for each 70^AG^ clone were normalized to signal from wildtype cells. The gating strategy is shown in Fig. S2B. Plot shows mean of data from three independent experiments, with SD. C: Ratio of 28S to 18S rRNA in WT and 70^AG^ cells, as determined by BioAnalyzer analysis. Mean and SD of data obtained in three independent experiments (total of 4 WT and 9 70AG samples).

The ratio between 18S and 28S did not vary significantly between wildtype and mutant cells in the FlowFISH data. To support the conclusion that 18S and 28S rRNA were equally affected, RNA samples were run on a BioAnalyzer to quantify the relative fluorescence between mature rRNA species (Fig. 3C)^32^. As expected, the ratio of 28S to 18S fluorescence was about 2 in all samples tested, with no significant differences between the mutants and parental cells.

As per-cell rRNA was reduced in 70^AG^ mutants, we next tested whether they also had a reduction in mature ribosomes using a recently developed technique called <u>T</u>otal <u>R</u>NA-<u>A</u>ssociated <u>P</u>rotein <u>P</u>urification (TRAPP)^19^. This method quantifies the RNA-bound proteome, with proteins recovered in proportion to their interaction with RNA. To do this, *in vivo* RNA:protein complexes are stabilized by UV irradiating living cells (Fig. 4A). Cells are then lysed in denaturing conditions, and RNA-associated proteins captured by binding the RNA portion of the complex to a silica column. Unbound proteins are washed away and the remaining silica-bound RNA-associated proteins digested with trypsin. Released peptides are eluted for analysis by mass spectrometry. Wildtype and 70^AG^ cells were directly compared by growing each in media with isotopically-labelled “heavy” or “light” amino acids, respectively, and mixing samples 1:1 by nucleic acid content. The output of the experiment is ratios of protein abundance between mutant and wildtype cells, called SILAC ratios (for Stable Isotope Labelling with Amino acids in Cell culture)^19,33^. We anticipated that the 70^AG^ mutants would show reduced recovery of ribosomal proteins (RPs) if mature ribosomes were less abundant in these cells.

**Figure 4:**
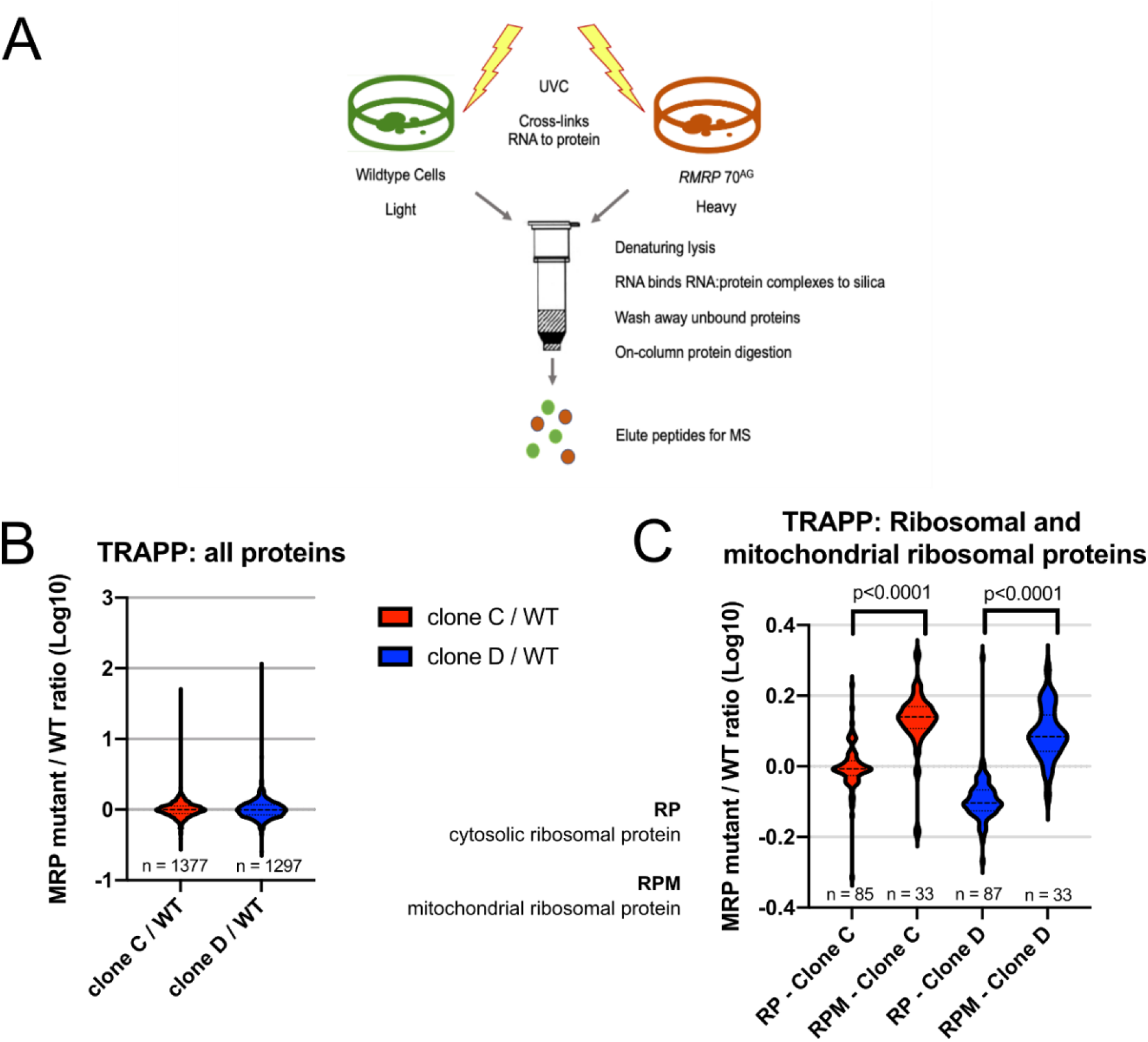
70^AG^ mutation in *RMRP* result in a reduced ratio of cytosolic to mitochondrial ribosomes. A: Overview of TRAPP technique. Wildtype and *RMRP* 70^AG^ cells were grown in heavy or light SILAC media, respectively, and UV crosslinked to stabilize RNA:protein interactions. Cells were lysed in denaturing conditions, and RNA:protein complexes bound to a silica column. Non-bound proteins were washed away, and remaining proteins digested on the column. Peptides were eluted for mass spectrometry. B: Log-transformed SILAC ratios (70^AG^ mutant/wildtype) for all proteins quantified in TRAPP. C: Log-transformed SILAC ratios for cytosolic ribosome proteins (RP) and mitochondrial ribosome proteins (RPM) obtained in TRAPP. Data shown in B and C are representative of two independent experiments, each including two SILAC mixes with different CRISPR-clones. Violin plots depict distribution of SILAC ratios, with lines at median and quartiles. Indicated p values derived from two-tailed t tests: for clone C, t=9.458, df=116; for clone D, t=13.62, df=118.

About 1,300 SILAC ratios were recovered per mix (Fig. 4B). The mean ratio was 1 (log transformed to 0), confirming that there was no systematic bias between samples (Fig. 4B). As expected, mutant cells showed a small decrease in average abundance of cytoplasmic RPs, implying reduced interaction between RPs and rRNA (Fig. 4C). Human cells have two separate populations of ribosomes, cytoplasmic and mitochondrial, which differ in both rRNA and protein composition^34^. Processing of mitochondrial ribosomes is independent of *RMRP*. Strikingly, recovery of cytoplasmic RPs from 70^AG^ cells was significantly lower than recovery of mitochondrial RPs, relative to wildtype cells (RPMs; Fig. 4C). This difference was highly statistically significant (p < 0.0001). The same result was obtained in a repeat experiment where SILAC labelling was swapped such that mutant cells were “heavy” and wildtype cells “light”, confirming that it was not a technical artifact caused by SILAC labelling.

There are two possible explanations for this result. First, 70^AG^ cells might genuinely have more mitochondrial ribosomes. RNase MRP was originally identified as cleaving an RNA primer required for mouse mitochondrial DNA replication. The 70^AG^ mutation could potentially increase the efficiency of this process and increase mitochondrial copy number^9^. Alternatively, the result could be caused by normalization to total RNA. In TRAPP, equal amounts of RNA from wildtype and mutant cells are mixed in order to purify RNA-bound proteins^19^. If mutant cells have less total RNA per cell, due to reduced rRNA abundance, more cell-equivalents will be included in the mix relative to the wild-type. This will cause an apparent reduction in ribosomal proteins in the TRAPP-purified proteome compared to other abundant RNA-interacting proteins, such as mitochondrial RPs.

Cytoplasmic RPs are also abundant relative to the total proteome so a similar effect would be expected for analyses total protein. Indeed, the same trend was seen in SILAC total proteome data (Figs. S4A and S4B).

### 70^AG^ mutation in *RMRP* reduces the abundance of intact RNase MRP complexes

RNase MRP is a ribonucleoprotein complex comprised of *RMRP* and about 10 proteins.^9^ All of these are reported to be shared with an evolutionarily related ribonucleoprotein complex, RNase P, which cleaves the 5’ leader from pre-tRNAs. RNase P includes the ncRNA *RPPH1* in place of *RMRP* ^9^ and has similar abundance to MRP^35,36^. Notably, mutations in the shared POP1 protein phenocopy the skeletal disease phenotype of *RMRP* mutations.^37-39^

In TRAPP data, the RNase MRP/P complex proteins POP1 and RPP38 showed consistently reduced RNA-association in 70^AG^ mutants compared to controls (Fig. 5A). In proteomic data, POP1 was slightly less abundant in mutants than parental cells, but this was not statistically significant (SD crossing 1; Fig. 5B). The results raised the question of which RNAs are most associated with POP1 *in vivo, as* these interactions must be reduced in 70^AG^ cells to cause the reduced recovery of POP1 in TRAPP. To address this, we used the technique of <u>cr</u>osslinking and <u>a</u>nalysis of <u>c</u>DNAs (CRAC) to map RNAs associated with individual MRP/P complex proteins^40^. At present there is no high-resolution structure of the human RNase MRP complex to inform choice of proteins for CRAC. We therefore selected three MRP/P proteins, POP1, POP4 and POP5, based on structural data from yeast suggesting these may be in close proximity to the known pre-rRNA substrate^41^. A CRISPR-based approach was used to insert a FLAG-HIS_7_ tag onto the N or C terminals of these proteins in K562 cells. Homozygous-tagged clones were selected for analysis and RNA:protein complexes were stabilized by UV crosslinking *in vivo*. The bait protein was then tandem purified in stringent, denaturing conditions, and co-purifying RNAs sequenced^40^.

**Figure 5:**
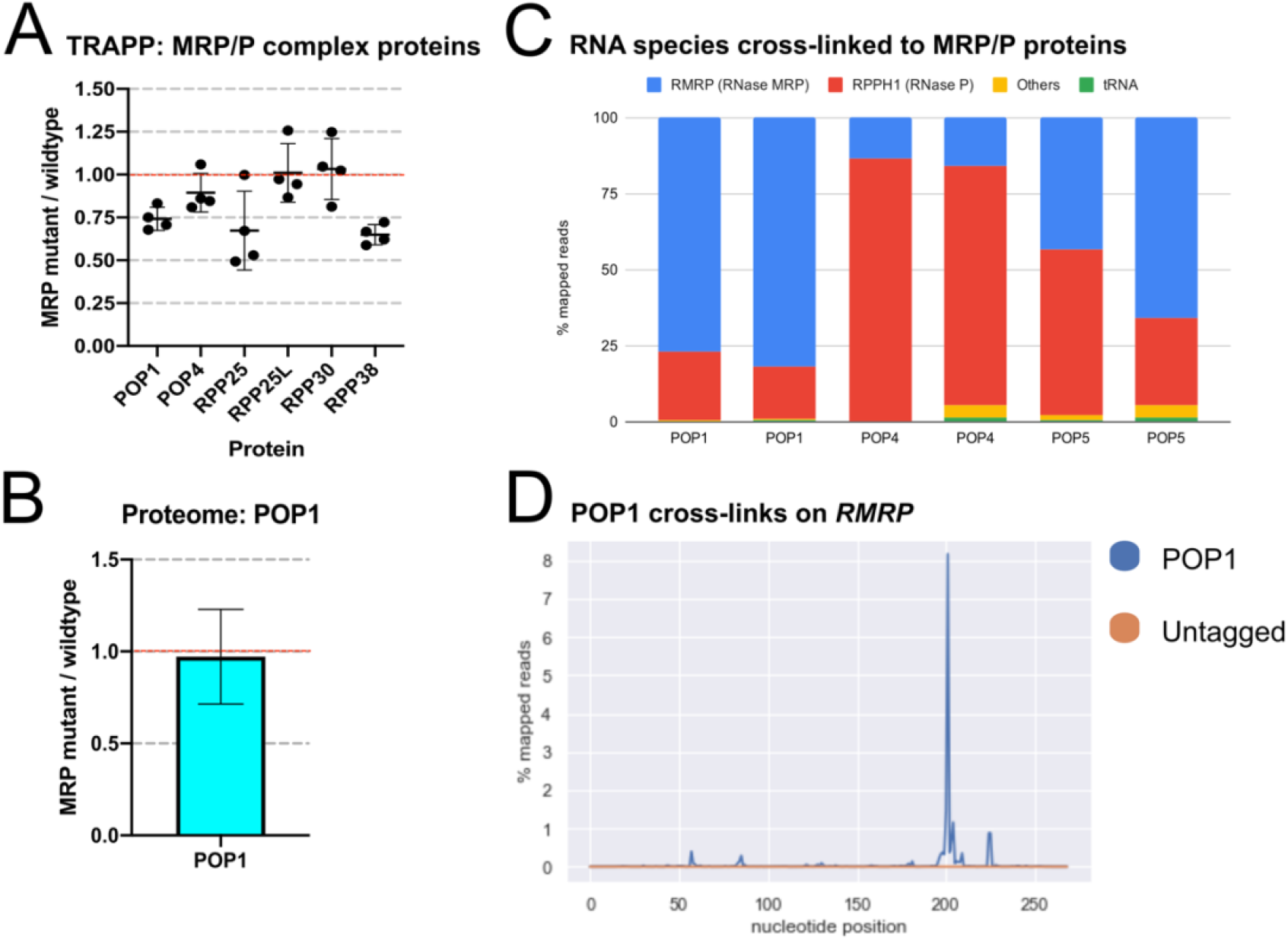
70^AG^ mutation in *RMRP* reduces the abundance of intact RNase MRP complexes. A: SILAC ratios (70^AG^ / wildtype cells) obtained for MR/P complex proteins in TRAPP (mean and SD). Each dot represents data from an independent SILAC mix. B: SILAC ratios (70^AG^ / wildtype cells) obtained for POP1 protein in proteome. Includes data from two SILAC mixes, showing mean and SD. C: RNA species recovered from MRP/P complex proteins in CRAC. Graphs shows relative proportion of mapped reads representing each indicated RNA or RNA biotype, in two independent experiments. D: Pileup of reads containing single base deletions (indicating RNA:protein crosslink site) in *RMRP*, in POP1 or negative control CRAC. Representative of results from two independent experiments.

Notably, *RMRP* comprised the majority of RNAs recovered with POP1, making up 77 - 82% of sequencing reads (Fig. 5C). *RPPH1* was significantly less recovered with POP1, accounting for 17 - 23% of reads. As the abundance of *RMRP* was not significantly reduced in 70^AG^ mutants (Fig. S3C), this result indicates that the reduced total RNA interactions of POP1 in 70^AG^ cells likely reflects impaired POP1:*RMRP* interactions. POP4 CRAC recovered the opposite ratio from POP1, with 79 – 86% of reads representing *RPPH1 (*Fig. 5C). In POP5 CRAC, the ratio of *RMRP* to *RPPH1* was variable between replicates, but approximately equal overall (43% vs 55%). Sites of RNA:protein crosslinks in CRAC are often revealed by single-base deletions in the sequencing data^40^. Mapping such crosslinks across *RMRP* showed a clear, single site of interaction with POP1, centered on nucleotide 201 (Fig. 5D), whereas crosslinks on *RPPH1* were more diffuse with several smaller peaks (Figs. S5A and S5B). Conversely, POP4 CRAC showed two clear peaks in *RPPH1* (at positions 122 and 186), but multiple smaller peaks for *RMRP* (Figs. S5C and S5D). We noted that the POP1 crosslinking site in *RMRP* is located just downstream of the eP19 domain, similar to the interaction site of yeast Pop1 on the NME1 RNA^36^. To determine whether the preference for POP1 crosslinking to the RNA component of MRP over P was also conserved, we tested yeast Pop1 in CRAC. This showed that, indeed, the bias was even more pronounced: 89 - 94% of reads were from the MRP RNA (Nme1 in yeast) and 3 - 7% from RNase P (Rpr1; Fig. S6).

We conclude that POP1 shows a conserved, preferential association with *RMRP* ncRNA *in vivo*. Levels of POP1 and *RMRP* are maintained in 70^AG^ cells, so the reduced POP1 recovery in TRAPP indicate that the mutation destabilizes the POP1:*RMRP* interaction. This is likely to reduce the abundance or activity of RNase MRP complexes.

## Discussion

Ribosomopathies are a diverse group of human disorders in which ribosome production or function is defective^42^. Clinical features of these disorders are variable and often tissue specific^43^. Bone marrow dysfunction leading to anemia or cytopenia is common, as are skeletal anomalies and an increased risk of cancer. Like CHH, many ribosomal disorders have surprisingly tissue-specific phenotypes, despite ribosomes being present in almost all cells^42^. Two broad hypotheses have been proposed to explain this. First, the phenotype may reflect decreased ribosome number or function in specific cell types that crucially depend on poorly translated mRNAs, vulnerable to reduced translation. In Diamond Blackfan anemia (DBA), in which mutations in ribosomal proteins cause defective erythropoiesis, ribosome numbers are reduced without altered composition^44^. This results in reduced translation of transcripts with unstructured 5’ untranslated regions (UTRs), including a specific reduction in the translation of GATA2, a key hematopoietic transcription factor. Alternatively, and not mutually exclusively, some cell types may rely on specialized ribosomes, the assembly of which may be affected by particular mutations^42^.

Here we present evidence that CHH is a ribosomopathy caused by a defect in pre-rRNA processing, the first such disorder to be described. Patient fibroblasts and a cell line with a disease-linked mutation (70^AG^ in *RMRP*) showed delayed rRNA processing, in a pattern consistent with decreased cleavage at the presumed RNase MRP target (site 2 in ITS1). These cells also have reduced rRNA per cell, and reduced intact cytosolic ribosomes relative to the mitochondrial ribosome pool. Moreover, the 70^AG^ mutation reduces the abundance of intact RNase MRP complexes, probably by destabilizing the interaction between *RMRP* and POP1. The overall effects on ribosome synthesis are modest, but this is expected from a disease-related mutation since carriers must develop almost normally in order to be classed as patients.

These results advance our understanding of CHH, but also raise some intriguing questions. Most notably, why are the phenotypes of DBA and CHH different, if reduced ribosome number is a common pathological mechanism? Some aspects of the two disorders do overlap; for example, some CHH patients have bone marrow dysfunction similar to that seen in DBA^45^. However, other aspects are unique, notably the T cell dysfunction which is life-limiting in CHH but not a feature of DBA^1^.

A high-resolution structure of the human RNase MRP complex would also aid understanding the pathophysiology of CHH. The results presented here could contribute to such a structure, by showing the main *in vivo* interaction site between POP1 and *RMRP*. Why POP1 preferentially interacts with *RMRP* over *RPPH1* across species is unknown, and will be an important question for further studies.

In conclusion, the results in this study point to CHH being a disorder of ribosome synthesis, and suggest experimental approaches to further explore this complex disease.

## Acknowledgments

We thank Prof. Sophie Hambleton from Newcastle University for providing patient and control fibroblasts, sourced from the Great North Biobank (REC number 16-NE-0002). We also thank Martin Waterfall and Fiona Rossi for technical assistance, and Sarah Walmsley and Edward Wallace for helpful discussions. This work was supported by Wellcome through an Edinburgh Clinical Academic Track fellowship to NR (213011), a Wellcome Principal Research Fellowship to DT (077248) and Wellcome Trust Investigator Award to RZ (WT205014/Z/16/Z). Work in the Wellcome Centre for Cell Biology is supported by a Centre Core grant (203149).

## Author contributions

NR, RZ and DT conceived the project and wrote the manuscript. NR, VS, DW, TT and CS performed experiments. NR, VS, DW, TT, AH, RZ and DT designed experiments. NR, VS, CS and TT analyzed data. All authors edited and reviewed the manuscript.

## Data availability

The GEO accession number for all sequence data reported in this paper is GSE171021. The proteomics data are available through the ProteomeXchange partner repository^46^ with the dataset identifier PXD025029.

## Code availability

Analysis scripts are available on request.

## Figures

**Figure S1:**
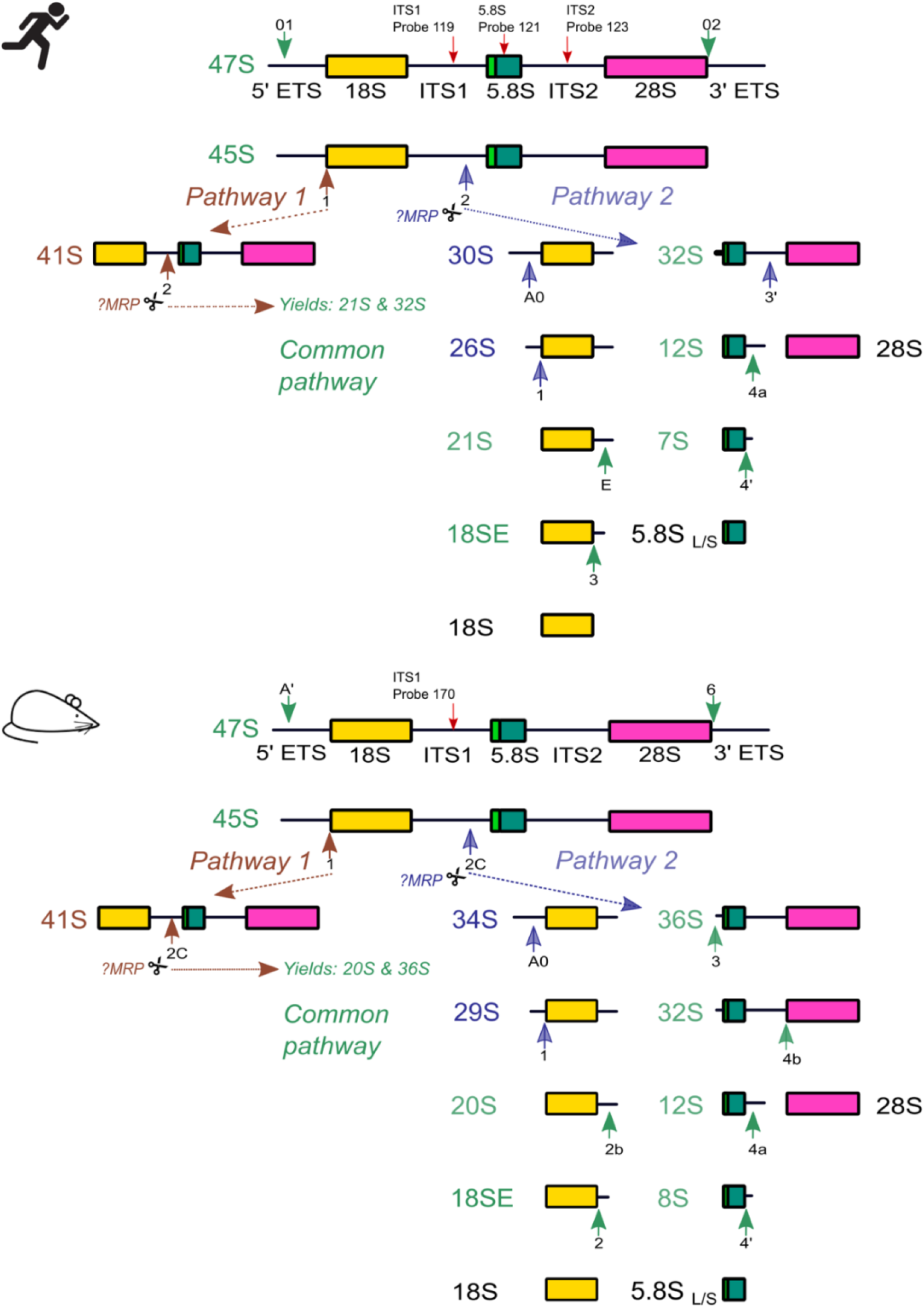
Pre-rRNA processing in humans and mice. Ribosomes are composed of ribosomal RNA (rRNA) and ribosomal proteins, and consist of a large subunit (LSU) and small subunit (SSU). 18S is the rRNA for the small ribosomal subunit. 5.8 and 28S are rRNAs in the large subunit. RNA encoding the mature 5.8S, 18S and 28S rRNA are transcribed together as a long precursor, called 47S in humans and mice. Both ends of 47S are flanked by external transcribed spacers (5’ ETS and 3’ ETS), and two internal transcribed spacers (ITS) separate the mature species. To make functional ribosomes, these spacers are removed in a complex series of processing steps. In humans, 47S is first processed by trimming the 5’ and 3’ ETS, by cleavage at sites 01 and 02, yielding the 45S precursor. After this, two distinct processing pathways have been described. Pathway 1 generates the 41S species by removing the remaining 5’ ETS, while the LSU and SSU precursors remain joined. In Pathway 2, the initial cleavage occurs at site 2 which separates the LSU and SSU. The most likely candidate for mediating this site 2 cleavage is RNase MRP^14^. The 41S species in Pathway 1 is also cleaved at site 2, yielding the 21S and 32S precursors which are then processed similarly to the same species generated in Pathway 2. In mice, pre-rRNA processing follows a similar pattern. The equivalent of site 2 is termed site 2C. In all studied fungi, animal and plants, 5.8S rRNA is present in both long and short forms, where the long form has a 7 to 8 nucleotide 5’ extension. The biological significance of the two forms is not known. The processing pathways depicted here are simplified for clarity, but show the major pre-rRNA species detected in experiments in this study. Solid green arrows indicate cleavage/processing at named sites in the common pathway. Brown and blue solid arrows represent cleavage/processing steps exclusive to Pathway 1 and Pathway 2, respectively. Dotted arrows indicate direction of processing. Red arrows indicate position of Northern probes used in this study. abundance of 41S to 47S/45S (PTP), and 30S to 41S, from Northern blots as illustrated in A.

**Figure S2:**
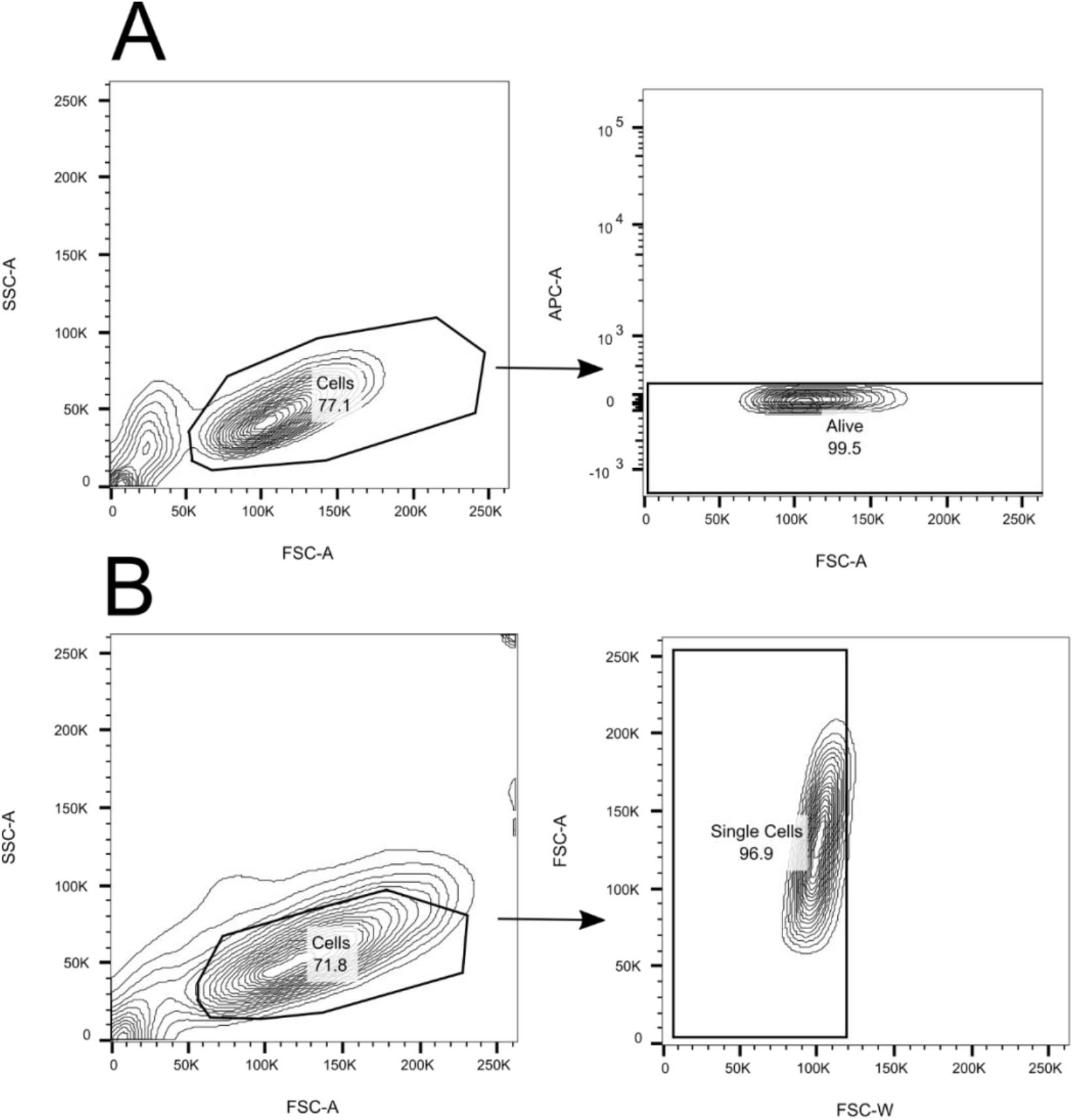
Gating strategies used in flow cytometry experiments. (A) Mouse T cell experiments: cells were gated first on morphological lymphocytes, then live cells as assessed by Zombie Red live-dead stain on the APC channel. (B) FlowFish experiments: K562 cells were gated first on morphologically live cells then on single cells.

**Figure S3:**
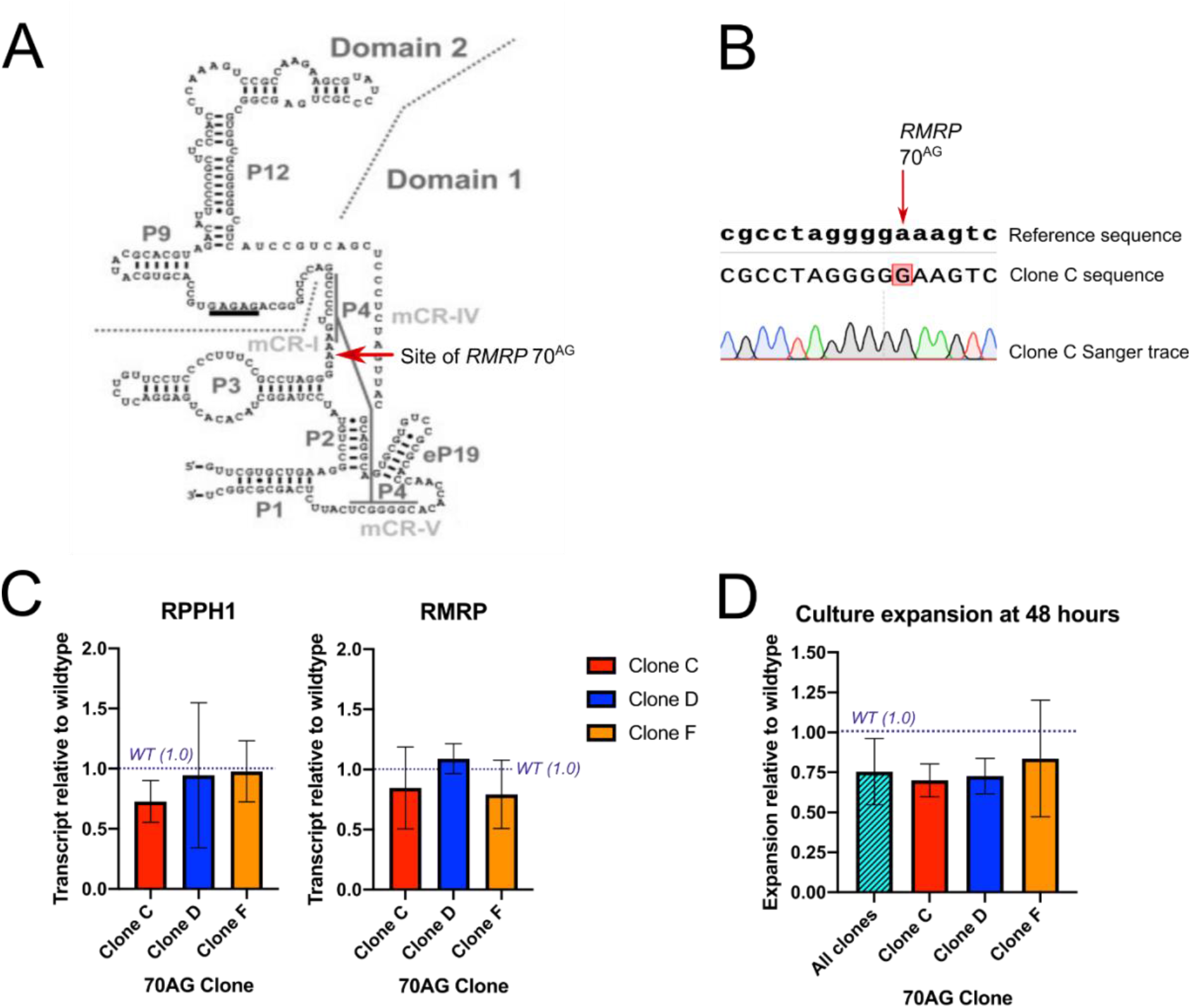
CRISPR-Cpf1 allows generation of K562 lines homozygous for *RMRP* 70^AG^ mutation. A: Predicted secondary structure of human *RMRP* showing site of 70^AG^ mutation. Adapted from Esakova and Krasilnikov.^36^ Regions evolutionarily conserved from M-type archaeal RNase P RNA are indicated as mCR-I, mCR-IV and mCR-V. Named base-paired RNA stems are indicted as P1, P2, etc. B: Extract from Sanger sequencing trace for *RMRP* 70^AG^ clone C. Similar results were obtained for the other clones used in this study. C: qPCR to quantify the RNase MRP ncRNA (*RMRP*) and the RNase P ncRNA (*RPPH1)* in *RMRP* 70^AG^ clones. Mean and SD deviation of results from three independent experiments, each normalized first to a house-keeping gene (*HPRT)* and then the average of wildtype values obtained in that experiment. D: Expansion of *RMRP* 70^AG^ and wildtype K562 cells after 48 hours of culture. Cells were plated at a density of 1 × 10^5^ / mL, in triplicate, and either counted manually every 24 hours, or tracked with live cell imaging software (Incucyte). Graph contains results obtained from one manual experiment and two live-cell imaging replicates (mean and SD).

**Figure S4:**
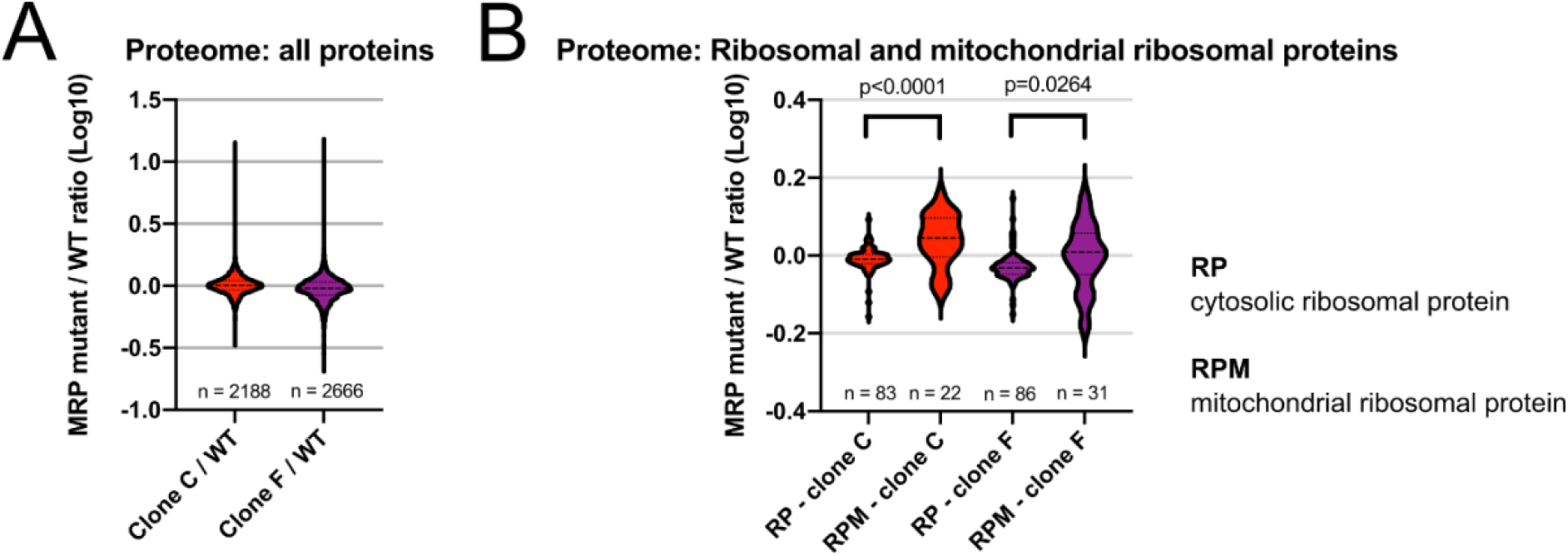
Proteome of *RMRP* 70^AG^ cells. A: Log-transformed SILAC ratios (70^AG^ mutant / wildtype cells) for all proteins quantified in two SILAC mixes using independent CRISPR clones. Lysates from indicated cells were grown in heavy or light SILAC media, mixed 1:1 by protein abundance and electrophoresed on a polyacrylamide gel. Proteins were digested ingel to release peptides for mass spectrometry. B: Log-transformed SILAC ratios for cytosolic ribosome proteins (RP) and mitochondrial ribosome proteins (RPM), obtained in two SILAC mixes. Violin plots depict distribution of ratios, with lines at median and quartiles. Indicated p-values derived from two-tailed t tests: for clone C, t=4.896,df=103; for clone F, t=2.249, df=115.

**Figure S5:**
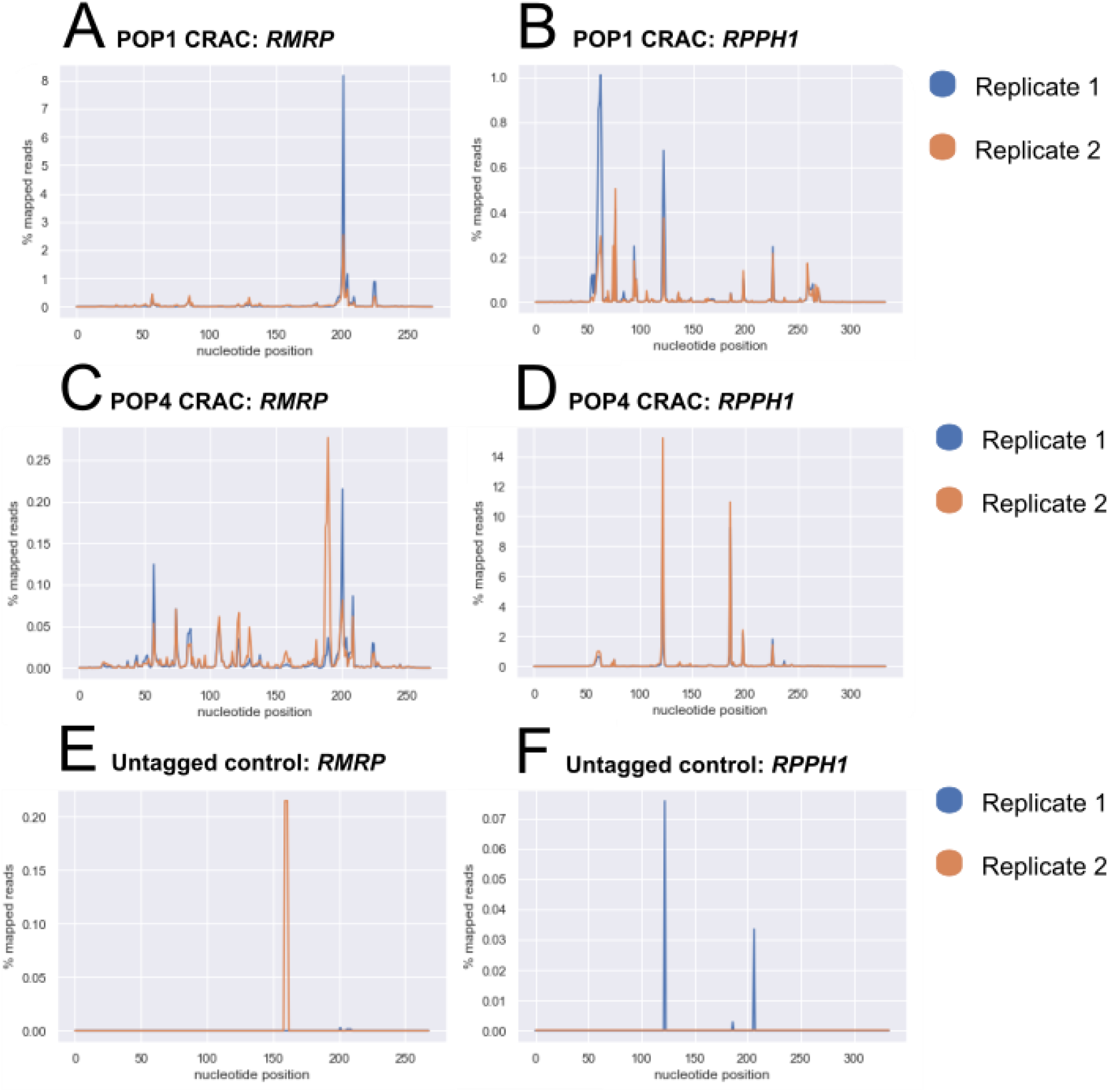
RNA:protein interactions in human MRP/P complexes. Sites of deletions, indicating RNA:protein crosslink sites, between human MRP/P complex proteins and ncRNAs. *RMRP* is the RNase MRP ncRNA. *RPPH1* is the RNase P ncRNA. A and B: Deletions in *RMRP* and *RPPH1* in POP1 CRAC. C and D: Deletions in *RMRP* and *RPPH1* in POP4 CRAC. E and F: Deletions in *RMRP* and *RPPH1* in negative control (untagged cells) CRAC. Graphs show proportion of mapped reads in two independent experiments. Note differences in scales used.

**Figure S6:**
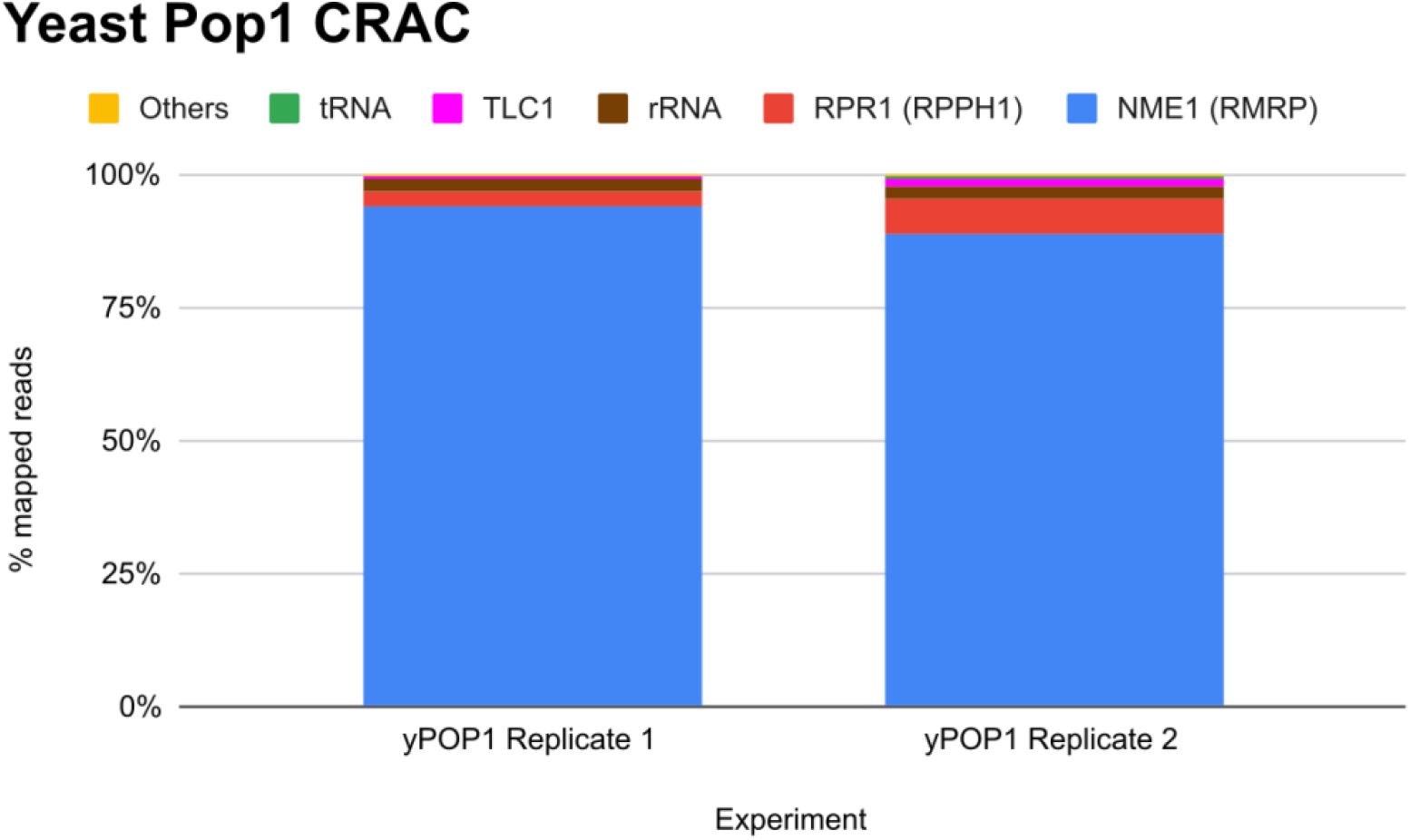
RNA species interacting with yeast Pop1 protein. RNA species and biotypes recovered in yeast Pop1 CRAC experiments. Graphs show relative proportions of mapped reads in two independent experiments. *TLC1* is the ncRNA component of the telomerase complex, a known Pop1 interactor in yeast^47^. *RPR1* is the yeast RNase P RNA, and *NME1* is the RNase MRP RNA.

## Supplementary Methods

### Human cell culture

K562 cells were obtained from ATCC (cat. CCL-243), and grown in RPMI 1640 Medium with GlutaMAX Supplement (Gibco; cat. 61870036), further supplemented with 1x final concentration Antibiotic-Antimycotic (Gibco; cat. 15240096) and 10% fetal calf serum (Sigma; cat. F2442). Cells were grown to a density of 0.5 - 1 × 10 ^6^ cells / mL, then diluted or used for experiments.

Patient and control fibroblasts were obtained from Prof. Sophie Hambleton (Newcastle University). They were grown in DMEM with GlutaMax (Gibco; cat. 10566016) supplemented with 10% fetal calf serum. Cells were grown to a confluency of 70 - 90% then split or used for experiments. To split, cells were washed once with sterile PBS, then incubated with 0.25% Trypsin-EDTA (Gibco; cat. 25200056; 0.2x volume of culture media removed) until detached. Trypsin was inactivated with 5 volumes of media, and cells pelleted at 100 RCF for 5 minutes before resuspension in culture media. All mammalian cell cultures were maintained at 37°C with 5% CO_2_

### CRISPR targeting of *RMRP* in primary mouse T cells

Rag1 KO, C57BL6/J mice were bred at the University of Edinburgh. All experimental procedures were approved under a project license granted by the UK Home Office, and additionally followed the University of Edinburgh’s ethical guidance.

Peripheral lymph nodes were dissected from Rag1 knockout mice homozygous for the OT1 allele. Lymph nodes were massaged through a 70 µM cell strainer. Cells were washed once with IMDM (Gibco; cat. 12440053) supplemented with 2mM L-glutamine, counted, and resuspended at 5 × 10^6^ cells/mL in PBS supplemented with 2.5 µM CellTrace Violet (Invitrogen; cat. C34557). After 20 minutes at 37 °C, cells were washed with complete T Cell media (TCM; IMDM supplemented with 10% FCS, 2 mM L-glutamine, 100 U/mL penicillin, 100 U/mL streptomycin, and 50 µM 2-mercaptoethanol), counted, and resuspended at 250,000 cells/mL in TCM supplemented with 10 nM N4 peptide (peptide sequence SIINFEKL). After 24 hours of stimulation, cells were counted, pelleted, and resuspended for transfection at 1.5 million cells per transfection in 80 µL Neon Transfection System Buffer T (Invitrogen; cat. MPK10096). CRISPR guide RNAs were purchased from IDT (Table 1), and resuspended at 100 µM in Nuclease-Free Duplex Buffer (IDT; cat. 11-01-03-01). For each transfection, 2.5 µL of guide RNA was mixed with 2.5 µL of tracrRNA and 20 µL of Nuclease-Free Duplex Buffer. The mix was heated at 95 °C for 5 minutes and then allowed to cool. Once at room temperature, 23 µL of Buffer T and 2 µL of TrueCut Cas9 Protein v2 (5 mg/mL stock; Invitrogen; cat A36496) was added, and the combined mix heated to 37 °C for 10 minutes before electroporation using a Neon Transfection System with 3x 1600 V pulses of width 10 ms. Electroporated cells were then cultured for 48 hours at either 250,000 (for flow cytometry) or 1 × 10^6^ cells per ml (for Northern blotting) in TCM supplemented with recombinant IL-2 (20 ng/mL). For flow cytometry, cells were first washed once with PBS before live/dead staining with Zombie Red Fixable Viability Kit (1:100 in PBS for 20 mins at room temperature; BioLegend; cat. 423109). Cells were then washed twice with FACS buffer before sample acquisition.

**Table 1:**
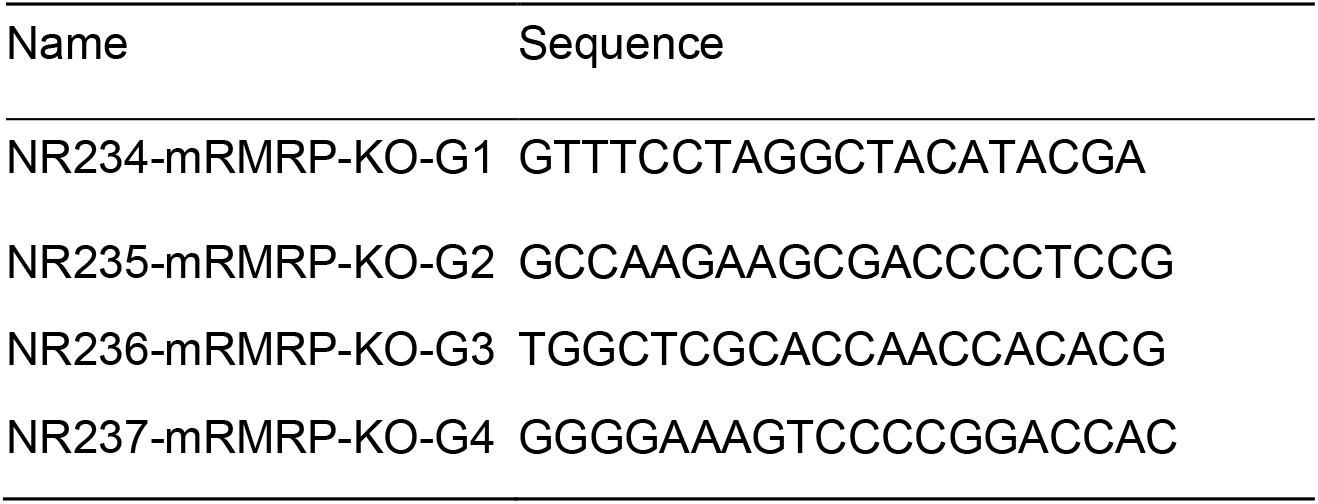
Guide RNAs used to target Rmrp in mouse T cells

### ICE-analysis of CRISPR-targeted mouse T cells

250,000 T cells were used as the input for DNA extraction, PCR and sequencing, using primers NR87 (5’-CCCACCTAGCGTTCCTACAT-3’) and NR88 (5’-AGAAATAAAAGTGGCCGGGC-3’). Sanger sequencing traces were analysed using the Synthego Inference of CRISPR Edits (ICE) tool web interface^16^.

### Flow Cytometry

For all flow cytometry assays, final washes and acquisition were done in FACS buffer, composed of 1x PBS supplemented with 2% FBS, 2 mM EDTA and 2 mM NaN_3_. Samples were acquired on a BD LSRFortessa, and samples exported to FlowJo 10.6 for analysis.

In all experiments, cells were first gated on morphological cells. For mouse CRISPR experiments, cells were further gated on live cells as determined by Zombie Red staining. For proliferation experiments, fold expansion of culture was calculated by summing all the cells in this gate, and dividing this by the calculated number of initial cells. Initial cell number was calculated by first calculating the number of cells in each division peak, and dividing this by 2^x^, where × is the number of divisions undergone by cells in that peak. Then, this value for all peaks was summed, giving the final initial cell value.

### Northern Blotting

#### RNA extraction

Mammalian cells were lysed in Trizol (Invitrogen; cat. 15596026). 200 µL of chloroform was added per 1 mL of Trizol used for cell lysis, and the mixture incubated at room temperature for 5 minutes. Phase separation, RNA precipitation and washes were done according to manufacturer’s protocol. After the wash step, the RNA pellet was resuspended in 100% formamide for Northern blotting, or water for qPCR. To ensure the RNA was fully dissolved, the mixture was left on ice for 20 minutes followed by heating at 65 °C for 10 minutes. A 1:10 dilution of RNA mix was quantified on a spectrophotometer.

#### Acrylamide gel for short RNA species

Loading buffer was made by supplementing 95% formamide with 20mM EDTA, made up with MQ. One crumb of Xylene cyanole and bromphenol blue was added per 5 mL of this buffer. RNA samples were mixed 1:1 with loading buffer and heated to 65 °C for 5 minutes, and then incubated on ice before loading.

Gel mix containing 8.3 M urea was prepared by mixing 50 mL of 40% acylamide gel solution (bis-acrylamide ratio 19:1; Severn Biotech; cat. 20-2400), 125 g of urea and 25 mL of 10x TBE buffer, and diluting to 250 mL with MQ. Before pouring, 300 µL of 10% ammonium persulfate and 30 µL of TEMED (Sigma; cat. GE17-1312-01) were added per 50 mL of gel mix.

Gels were run for 1600 V hours (for example, 80 V for 20 hours), and samples transferred onto BrightStar-Plus Positively Charged Nylon Membrane (Invitrogen; cat. AM10100) in 0.5x TBE for 3 hours at 40 V. The membrane was then cross-linked at 254 nM in a Statralinker cross-linking device (Stratagene).

#### Agarose gel for long RNA species

50x TRI/TRI buffer was prepared by mixing 10 mL of triethanolamine and 13.5 g tricine, and bringing the volume to 50 mL with MQ. Agarose gel mix was prepared by combining 285 mL of MQ, 3 g of agarose and 6 mL of 50x TRI/TRI. Agarose was melted in a microwave and allowed to cool to just above room temperature, at which point 10.5 mL of 36% formaldehyde was added and the gel poured.

Gel loading dye was prepared by mixing 84 µL of 50x TRI/TRI, 4 µL of 0.5 M EDTA, 80 µL of 1% Bromophenol Blue and 1.84 mL of MQ. Pre-mix was then prepared by mixing loading dye, 36% formaldehyde and 1 mg/mL ethidium bromide in the ratio 14:1:1. RNA samples were mixed with an equal volume of pre-mix, and heated at 70 °C for 10 minutes, followed by cooling on ice for 5 minutes before loading. The gel was run at 140 V for 4.5 hours. After visualisation, RNA was transferred to BrightStar-Plus Positively Charged Nylon Membrane by overnight downward capillary transfer in 10X SSC. Membranes were then cross-linked, as above.

#### Radioactive olionucleotide probing of Northern membranes

100x Denhardt’s solution was prepared by dissolving 1 g of BSA fraction V, 1 g of Ficoll 400 and 1 g of PVP in 50 mL of MQ. Hybridisation solution was prepared by combining 25 mL of 20x SSC, 5 mL of 10% SDS and 5 mL of 100x Denhardt’s solution, and bringing to 100 mL with MQ. Oligo labelling reaction mix was prepared by mixing 1 µL of 10 mM oligo with 13 µL of MQ, 2 µL of 10x T4 PNK Reaction Buffer (NEB; cat. B0201S), 1 µL of 100 mM DTT, 1 µL of T4 PNK (NEB; cat. M0201L) and 3 µL of ^32^P-*γ*ATP (10 µCi/ µL; Hartmann Analytic). The labelling reaction was performed at 37 °C for 1 hour. In the meantime, the membrane was pre-hybridised in hybridisation solution at 50 °C with shaking. Following labelling, unincorporated ATP was removed from the reaction mix using a mini-Quick Spin Oligo Column (Roche; cat. 1814397001), following provided instructions. Recovered reaction mix was added to the membrane in 100 mL of hybridisation solution for overnight hybridisation.

The following morning, the membrane was washed four times with hybridization wash solution (2× SSC, 0.1% SDS). Two washes were done at room temperature, followed by a wash at 50 °C before a final room temperature wash. Membranes were then exposed to a phosphoimager screen. Before re-hyrbidisation with a different probe, membranes were stripped by incubating twice for 10 minutes with boiling stripping solution (0.1× SSC, 0.1% SDS). Sequences of oligonucleotide probes used are shown in Table 2.

**Table 2:**
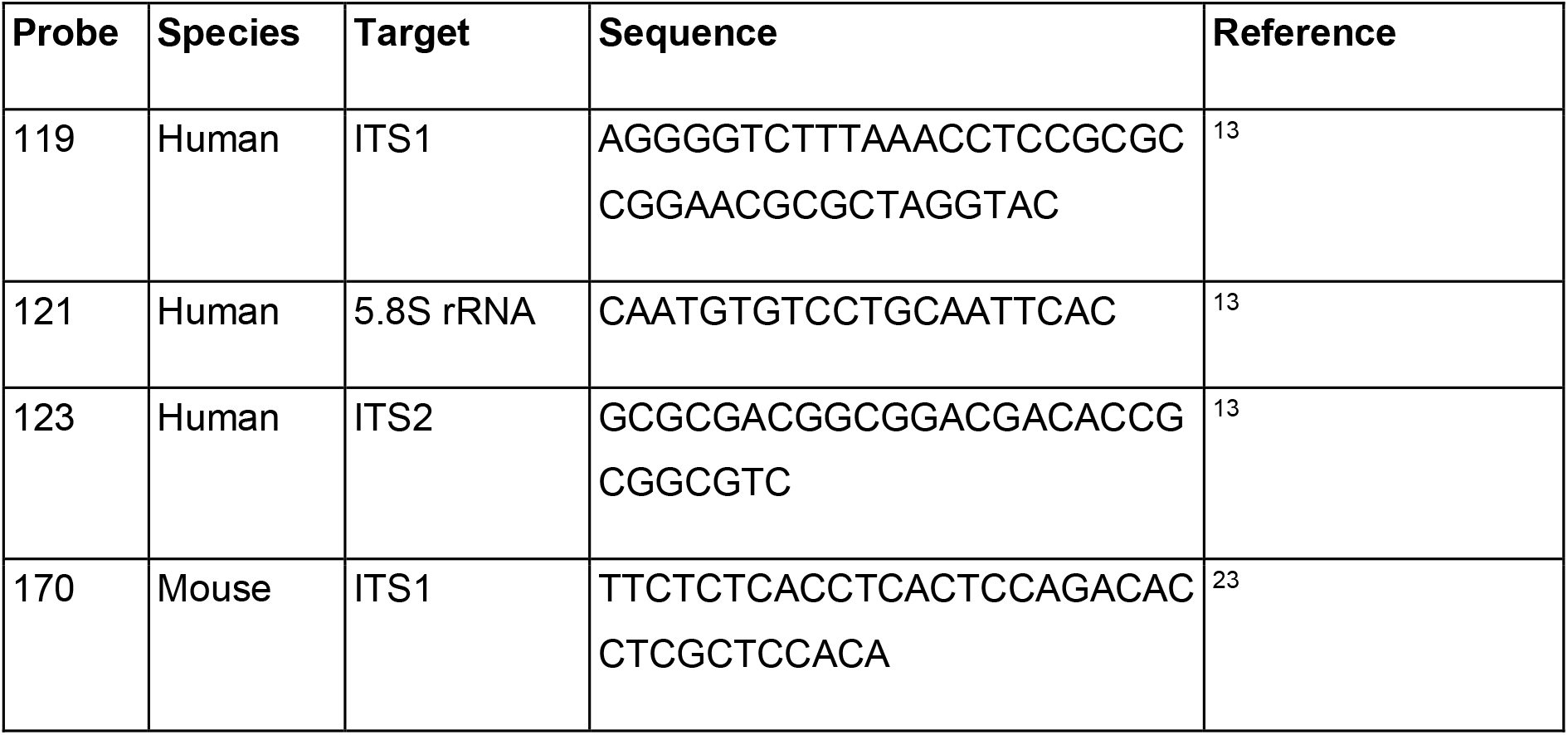
Oligonucleotide probes used for mammalian Northern Blotting experiments

### Generation of stable CRISPR-edited human cell lines

For all human CRISPR experiments, guide RNAs, repair templates and check primers were ordered from IDT from the sequences in Table 3.

**Table 3:**
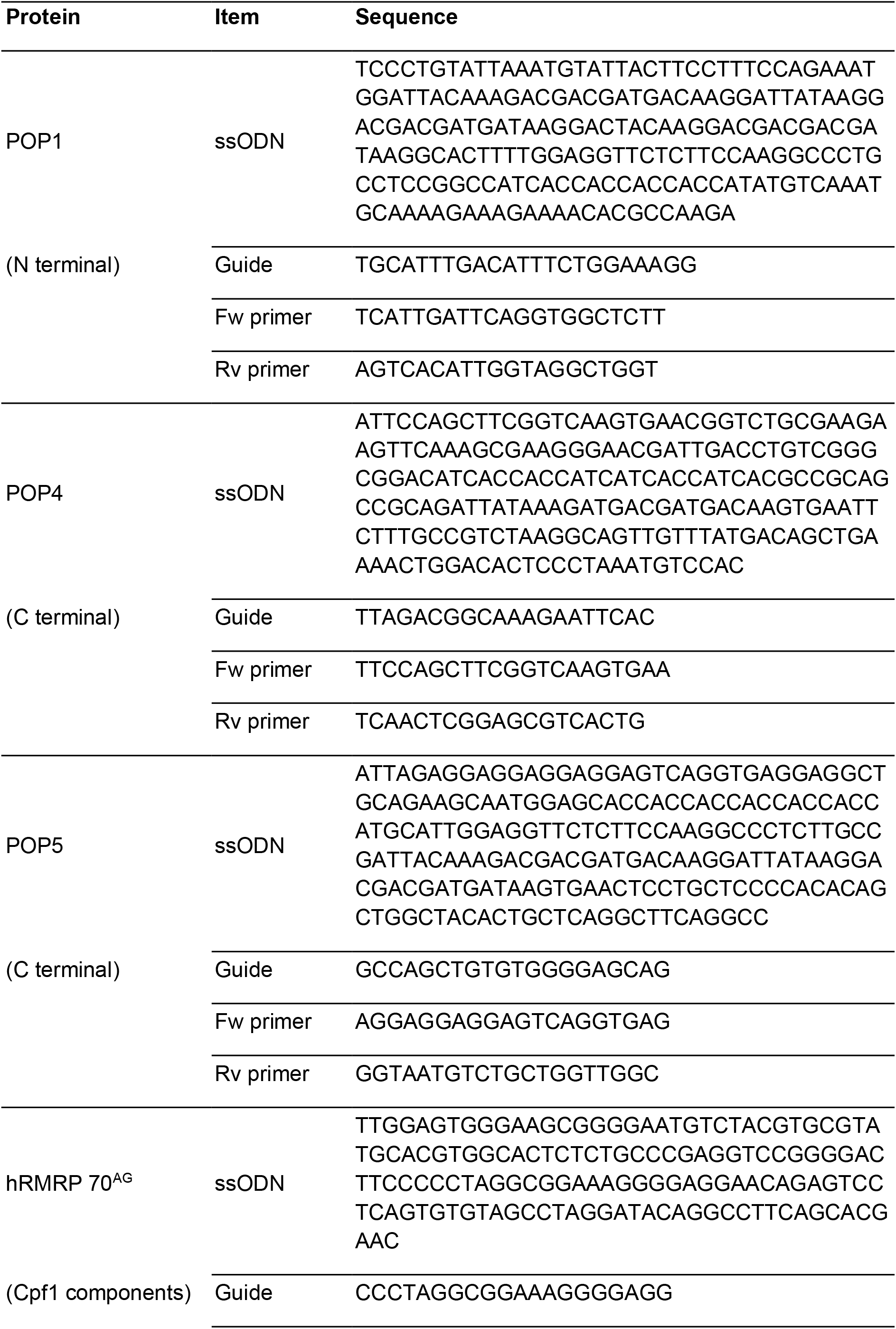
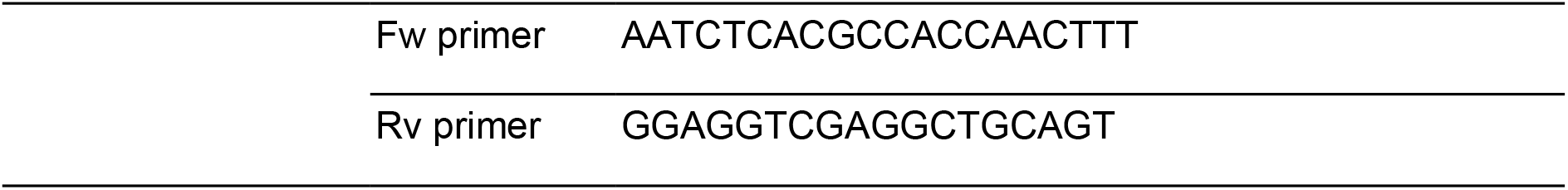
Components used for CRISPR-mediated editing of human cells

#### Protein tagging in human cell lines

For protein tagging experiments, single-stranded templates were designed including the tag with homology arms. For POP1, a 2xFLAG-6xHIS tag was inserted at the N terminus. For POP5, a C-terminal 2xFLAG-7xHIS tag was used. In the POP1 and POP5 tags, the FLAG and HIS moieties were separated a PreScission protease target sequence (amino acid sequence LEVLFQ/GP), but this was not used in the final purification protocol. For POP4, a C terminal 8xHIS 1xFLAG tag was used, with the HIS and FLAG moieties separated by a 4x Ala linker.

To introduce the tags, 0.6 µL of 200 µM guide RNA was mixed with 0.6 µL of 200 µM tracrRNA (Alt-R CRISPR-Cas9 tracrRNA (IDT; cat. 1072532), heated to 95 °C for 5 minutes then allowed to cool. Then, 2.1 µL of PBS and 1.7 µL of 61 µM Cas9 enzyme was added (Alt-R S.p. Cas9 Nuclease V3; IDT; cat. 1081058) and the mixture incubated at room temperature for 20 minutes. Meanwhile, 1 × 10^6^ cells per transfection were pelleted, washed with PBS and resuspended in 100 µL of Nucleofector Solution from the Cell Line Nucleofector Kit V (Lonza; cat; VVCA-1003).

Transfection mix was assembled with 5 µL of Cas9:gRNA mix (made above), 1 µL of 100 µM electroporation enhancer (IDT; cat.1075915) and 3 µL of 10 µM ssODN. Cells and transfection mix were combined and electroporated with a Lonza Nucleofector 2b Device, using supplied settings for K562 cells. Cells were then gently transferred to 1.5 mL of K562 media supplemented with 25 µM HDR Enhancer (IDT; cat. 1081072). After 48 hours, cells were sorted by FACS to 1 cell / well in a 96 well plate. Clones were screened by PCR and sequencing after about two weeks.

#### CRISPR-Cpf1 mediated knock-in of RMRP 70^AG^ mutation

A single-stranded repair template was designed including the mutation with homology arms. 1.6 µL of 100 µM Cpf1 crRNA was mixed with 1.4 µL of PBS and 2 µL of Cpf1 Nuclease 2NLS (IDT; discontinued) and incubated for 20 minutes at RT. Thereafter the procedure was the same as for CRISPR-Cas9 mediated protein tagging, except the electroporation enhancer was Cpf1-specific (IDT; cat. 1076300).

### qPCR

RNA was extracted from 1 × 10^6^ cells, and 5 µg of RNA diluted to 10 µL in MQ. Reverse primer mix was made by combining 15 µL of all the reverse primers listed in Table 4, each at 100 µM. 2.5 µL of this mix was added to each 10 µL RNA sample, and the combined mix heated to 72 °C for 5 minutes, after which the mix was split into two: one portion for reverse transcription and the other for a no-reverse transcription control.

**Table 4:**
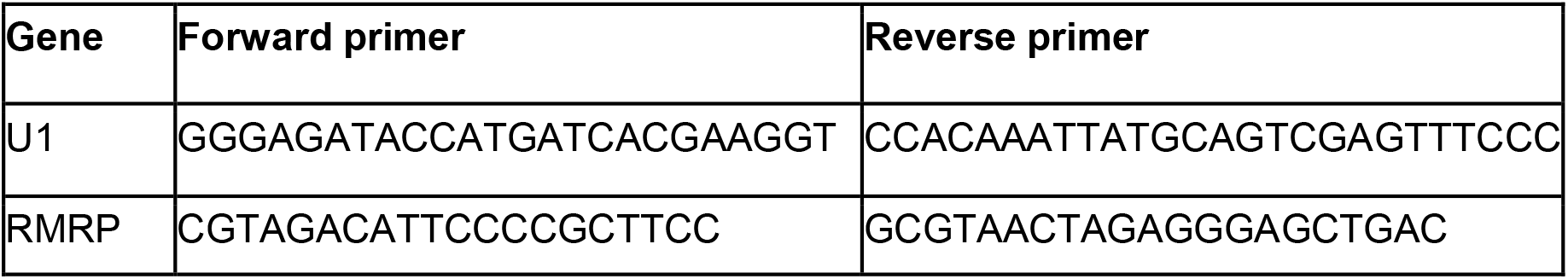
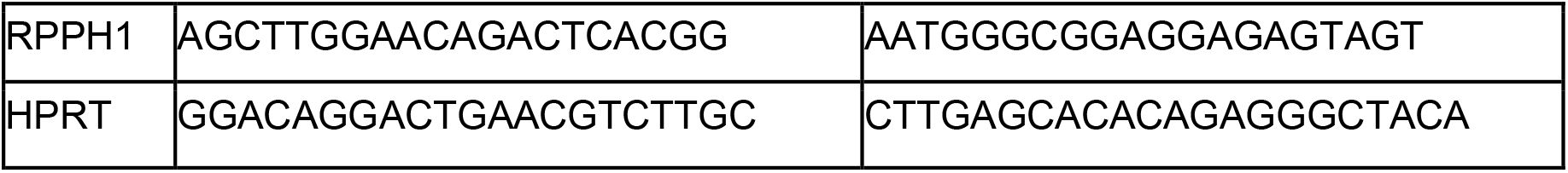
Primers used for qPCR experiments on human cell lines

Reverse transcription mix was made by combining 1.75 µL of MQ, 0.75 µL of 10 mM dNTP solution, 0.1 µL RNasin Ribonuclease Inhibitor (Promega; cat. N211A), 0.4 µL of Superscript IV Reverse Transcriptase (Invitogen; cat 18090010) and 2 µL of provided reaction buffer. Mix and RNA samples were combined, incubated at 55 °C for 1 hour, then diluted 1:200 for use.

qPCR reaction mix was made by combining 2 µL of cDNA or no-reverse transcription control, 0.1 µL of MQ, 0.2 each of forward and reverse primers (at 10 µM; Table 4) and 2.5 µL 2x SYBR Green PCR Master Mix (Applied Biosystems; cat. 4344463). Each reaction was set up in triplicate. cDNA was amplified in a LightCycler 480 (Roche), using the following cycle: initial 5 minutes at 95 °C, then 40 cycles of 10 s at 94 °C, 10 s at 60 °C and 15 s at 72 °C.

C_T_ value for each amplification curve was determined by the LightCycler software, and averaged for technical triplicates. Matched C_T_ for the house-keeping gene was subtracted to give ΔC_T_. Fold change of mutant (Mut) cells compared to wildtype (Wt) cells was calculated with the formula: 2 - ^(ΔCT Mut - ΔCT Wt).^

### FlowFISH

The FlowFISH method^17^ was adapted. Probe sequences shown in Table 5 are published^48^. Probes conjugated to fluorophores were ordered from IDT.

**Table 5:**
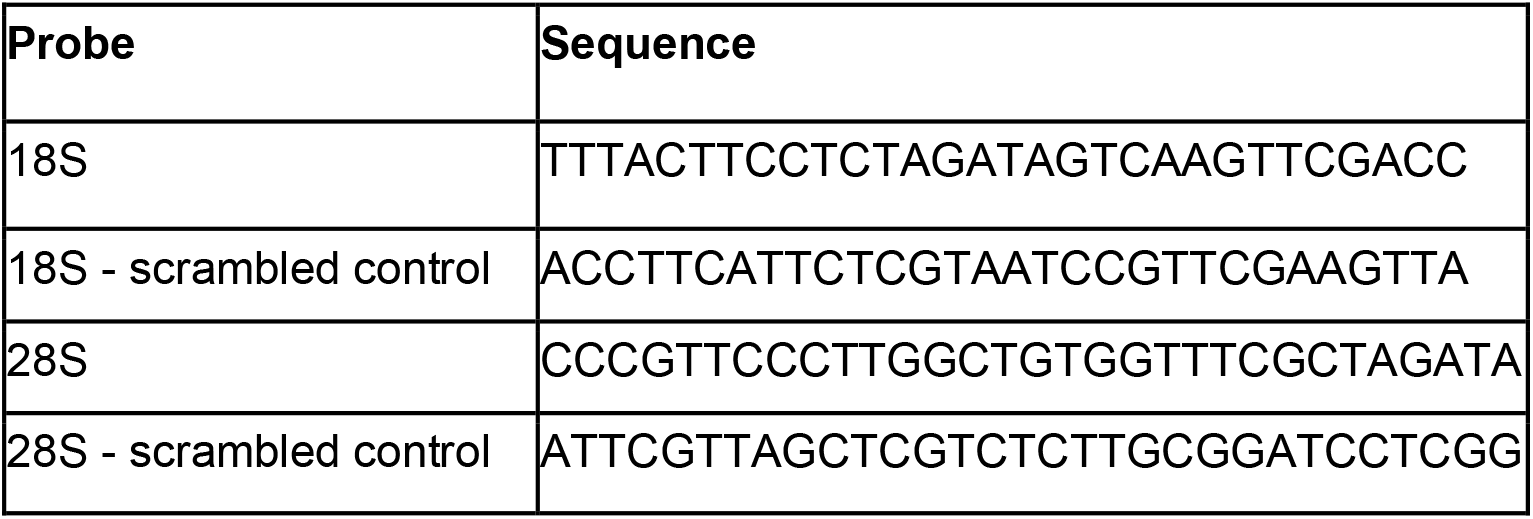
rRNA probes used for FlowFISH experiments

#### Fixation and permeabilization

0.5 × 10^6^ K562 cells were used. Cells were pelleted at 500 g for 5 minutes at room temperature, and washed once with 0.5 mL of PBS. After pelleting, cells were resuspended in 0.5 mL of PBS, and 0.5 mL of 8% paraformaldehyde added. Cells were left to fix for 30 minutes at room temperature, after which they were washed twice with 1 mL of PBS and resuspended in 0.5 mL PBS. 0.5 mL of 70% ethanol was then added dropwise, and cells pelleted again before resuspension in 1 mL of 70% ethanol and permeabilization overnight at 4°C.

#### Rehydration and probing

FlowFISH wash buffer (FFWB) was prepared by supplementing 2x SSC with 10% formamide and 0.25 mg/mL Bovine Serum Albumin fraction V (BSA; Sigma; cat. 05482).

FlowFISH solution A (FFSA) was prepared by combining, per sample: 5 µL of formamide; 2.5 µL of 2x SSC; 2.5 µL of 10 mg/mL *E. coli* tRNA (Sigma; cat. R1753); 2.5 µL of FISH probes diluted to 50 ng/µL; and 8.75 µL of MQ. FFSA was then heated to 95 °C and allowed to cool. Meanwhile, FlowFISH solution B (FFSB) was prepared by combining, per sample: 25 µL of 20% dextran sulphate dissolved in 4x SSC; 1.25 µL of 10 mg / mL BSA; and 40 units of RNasin Ribonuclease Inhibitor (Promega; cat. N211A). Once FFSA was cool, FFSA and FFSB were combined 1:1 to create the staining mix.

For staining, cells in ethanol were pelleted at 1000 g for 5 minutes, resuspended in 1 mL of FFWB and left to rehydrate at room temperature for 5 minutes before again being pelleted. Cells were then resuspended in staining mix for 3 hours at 37°C, before 2 washes with 1 mL FF wash buffer, and 2 washes with 1 mL FACS buffer.

### Total RNA-Associated Protein Purification (TRAPP)

K562 were grown for 10 divisions in SILAC RPMI (Thermo Fischer; cat. 88365) supplemented with 10% dialysed FBS (Gibco; cat. 26400044) and 50 µg/L each of lysine and arginine. For “light” cultures, these amino acids were obtained from Sigma. For “heavy” cultures, ^13^C_6_-lysine and ^13^C_6_-arginine were obtained from Cambridge Isotope Laboratories (cat. CLM-226 and CLM-2247, respectively).

Cells were grown to a density of 0.5 - 0.8 × 10^6^ cells / mL, with minimum 90% viability as determined by Trypan blue exclusion (Invitrogen; cat. T10282) using a Countess automated cell counter. 25 mL aliquots of culture were transferred to a custom quartz dish, and cross-linked with 400 mJ/cm^2^ of UVC using a Vari-X-Link device.^18^ Enough aliquots were cross-linked to create a sample containing 50 × 10^6^ cells. After cross-linking, cells were pelleted, washed with ice-cold PBS, and frozen at -80 °C.

RNA-associated proteins were purified following a modified version of the published TRAPP protocol, where silica columns were used in place of silica beads^19^. Proteins were digested on the column with 0.25 µg of Trypsin/Lys-C protease mix (Promega; cat. V5071), and peptides eluted for mass spectrometry. Samples were acquired by the Proteomics Facility at the Wellcome Centre for Cell Biology, University of Edinburgh, on an Orbitrap Fusion Lumos Tribrid Mass Spectrometer (Thermo Fisher Scientific, UK). Raw data was processed by the MaxQuant software platform, version 1.6.1, searching against the UniProt reference proteome set. Further analysis used custom scripts. A paper describing this modified TRAPP protocol is in preparation.

### HIS-TEV-Protein A tagging of yeast Pop1

Yeast stains in this study were derived from *Saccharomyces cerevisiae* strain BY4741^49^. HIS-TEV-Protein A (HTP) inserts with homology arms were amplified from plasmid pBS1539 using primers listed in Table 6. The transformation protocol described here is adapted from Gietz and Woods^50^. An overnight yeast culture was diluted to 0.5 × 10^7^ cells / mL (5 mL per transformation) and grown for 2 divisions. Cells were pelleted by centrifuging at 3000 RPM for 2 minutes at room temperature, washed with MQ, and resuspended in transformation mix composed of 240 µL PEG 3350 50% w/v, 34 µL PCR product, 50 µL pre-boiled salmon sperm DNA and 36 µL 1 M LiAc.

**Table 6:**
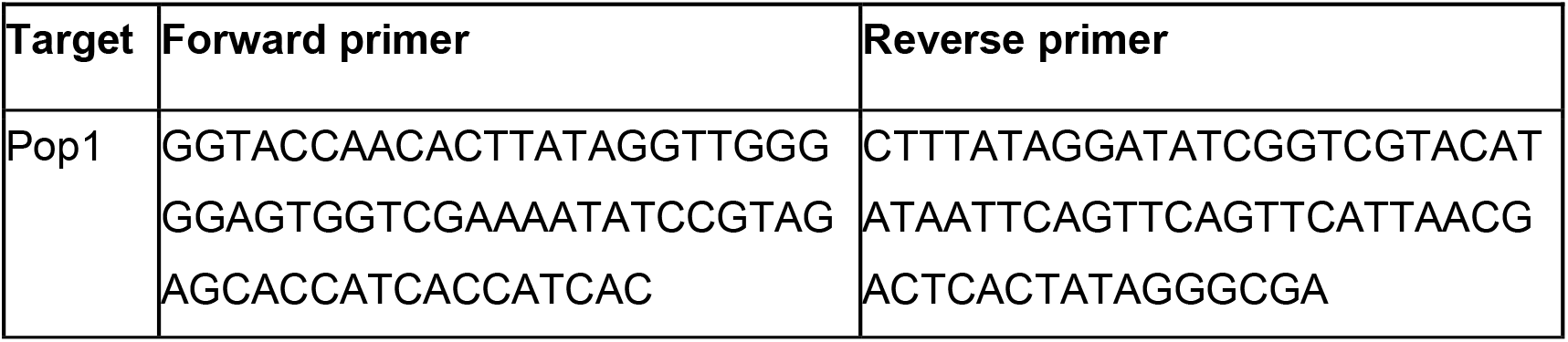
Primers used Pop1 tagging in yeast cells

The mix was vortexed and incubated for 40 minutes at 42 °C. Cells were then pelleted at 14,000 RPM for 30 s at room temperature, resuspended in 80 µL MQ, and spread on a YPD plate. After two days, colonies were streaked onto selective medium (-URA).

### Cross-linking and analysis of cDNAs (CRAC)

#### Preparation of human CRAC cells

Cells were grown and cross-linked as for TRAPP, except that normal RPMI media was used rather than SILAC media.

#### Preparation of samples for yeast Pop1 CRAC

2.75 L of yeast culture was grown to 0.5 OD, and irradiated for 100 s with UVC (254 nM) in a custom “Megatron” cross-linking device, as described^40^. Cells were then pelleted (2700 RCF for 15 minutes at 4 °C) and washed with ice-cold PBS, pelleted again and frozen at -80 °C until use.

#### CRAC protocol

For yeast Pop1, CRAC was performed as described^20^. For human CRAC, technical modifications were made as these cells had a FLAG tag in place of Protein A. anti-FLAG M2 magnetic beads (Milipore; cat. M8823) were used instead of IgG-coupled sepharose beads. To elute, anti-FLAG beads were resuspended in 200 µL of lysis buffer supplemented with 150 µg / mL of 3x FLAG peptide (Sigma; cat. F4799), and incubated for 5 minutes at 37 °C, shaking at 1200 RPM. The eluate was collected and the elution repeated. The two eluates were pooled, the GhCL added to denature proteins, as in the published protocol. Various other minor changes were made to buffer composition, lysis conditions and enzymatic steps. A paper describing this protocol is in preparation.

#### Sequencing and analysis of CRAC data

CRAC libraries were sequenced either on a MiniSeq or HiSeq, both with 150 base reads. Yeast CRAC data were processed using custom scripts calling utilities from the PyCRAC collection^21^. First, raw FASTQ files were demultiplexed using pyBarcodeFilter. Then, adapters and low-quality sequences were removed with Flexbar (version. 3.4.0)^51^. Next, PCR duplicates were collapsed with pyFastqDuplicateRemover. Reads were then aligned to the *Saccharomyces cerevisiae* genome sequence (Ensembl release EF4.74) by Novoalign version 2.07.00. Read counts were produced using pyReadCounters, and pileups with pyPileup. Graphs were produced with custom Python3 scripts, available on request. For human data, low quality reads were removed with Flexbar. STAR (version. 2.7.3a) was then used to align reads to a custom transcriptome database, based on GRCh38 with additional tRNA and rRNA species included. Read counts and pileups were generated with custom scripts, available on request.

### Proteomics

Cells were lysed in 1x Laemmli buffer, quantified and boiled for 5 minutes. 20 µg of protein was run per lane on a 12% pre-cast gel (BioRad; cat 4561043), with size determined using a pre-stained protein ladder run in lane 1 (NEB; cat. P7719S). Gel was washed three times with MQ, each for 5 minutes with nutation, then stained with Imperial protein stain (Thermo Scientific; cat. 24615) for 1 hour, and destained with MQ overnight. The gel was cut into fractions, and each fraction dissected into cubes not more than 1 mm^3^ in volume. Proteins were then digested with trypsin and StageTips created, following a published protocol^*52*^. Samples were acquired and analysed as described for TRAPP.

